# Uncovering the simple adhesive strategy of the Toxoplasma parasite for high-speed motility

**DOI:** 10.1101/2024.02.13.580110

**Authors:** Luis Vigetti, Bastien Touquet, Delphine Debarre, Thierry Rose, Lionel Bureau, Dima Abdallah, Galina V. Dubacheva, Isabelle Tardieux

**Author notes:** The authors contributed equally to this work (Luis Vigetti, Bastien Touquet).

## Abstract

*Toxoplasma gondii* is a protozoan parasite that has evolved a developmental morphotype called tachyzoite that navigates between cells and moves in and out of them in a wide repertoire of homeothermic hosts. Relying on a uniquely constant apicobasal bipolarity coupled to an actomyosin-driven retrograde surface flow, the tachyzoite has elaborated a molecular machinery to assemble transient anchoring contacts with the environment, which support the traction force required to power a typical helical gliding motility. Combining micropatterning with live, reflection interference contrast and expansion microscopies, we bring first nanoscale evidence that the tachyzoite needs to build only one apical anchoring contact with the substrate, thus spatially defining a minimal force transmission platform over which it can slide. We uncover that the apicobasal driven surface flow is set up in response to extracellular biochemical cues independent of adhesin release and tachyzoite-surface interactions, hence prior to motile activity. Furthermore, to identify the minimal adhesion requirements for helical gliding at the level of individual molecular species, we combine biochemical and biophysical quantitative assays based on tunable surface chemistry and quartz crystal microbalance with dissipation monitoring. These approaches uncover that glycosaminoglycan (GAG)-parasite interactions are sufficient to promote a productive contact for helical gliding and pave the way for the characterization of the structure and density of the molecules functionally engaged at this essential parasite-substrate mechanosensitive interface.

## INTRODUCTION

Cell locomotion achieved by eukaryotic individual cells and collectives has been phenomenologically categorized in a spectrum of modes from crawling or mesenchymal to amoeboid-like or bleb-based movements (**Extended Fig. 1a**). Although the actomyosin contractile system is generally involved in all modes, it can generate force through different architectures, and each of them in turn has to be combined with specialized adhesion structures, which also depend on the microenvironment (Bershadsky & Kozlov, 2011; Paluch et al., 2016). Furthermore, many protozoan and metazoan cells adjust to the fluctuating biochemical and biophysical environmental cues by switching from one mode to the others (Lämmermann & Sixt, 2009; Liu et al., 2015; Yamada & Sixt, 2019). Indeed, there is a renewed interest for theoretical and experimental studies on how eukaryotic cells integrate extracellular and intracellular variable inputs to tune their interactions with the extracellular matrices (ECMs) and adapt their migration strategies to fulfill specialized functions.

In this context, the protozoan *Toxoplasma gondii* is an experimentally accessible model system to interrogate the spatiotemporal coupling between cell-substrate adhesion and the contractile machinery required for cell movement. *T. gondii* is one of the most prevalent parasitic microbes of warm-blooded animals which might colonize about a third of the human population (Montoya & Liesenfeld, 2004). It is the etiological agent of toxoplasmosis, a set of potentially life-threatening or debilitating pathologies mostly in immunocompromised hosts (Milne et al., 2020). Tissue damages are primarily caused by the motile and fast-cycling developmental morphotype called tachyzoite that locomotes between cells in various ECMs and moves in and out of host cells in which it produces progeny (Frénal et al., 2017; Hu et al., 2002; Leung et al., 2014; Pavlou et al., 2019). Using a peculiar adhesion-dependent motility mode referred as to gliding, the tachyzoite displaces its roughly seven micron-long body in just a handful of seconds, which is an impressive achievement for non-swimming eukaryotic cells (Philips & Milo, 2015).

The main simplifying feature of the tachyzoite model comes from the permanent apicobasal cell polarity, which defines one constant symmetry axis as early as biogenesis starts within the mother cell (Hu et al., 2002). In particular, the apical pole is specified by a conical tubulin-based appendage known as conoid, which houses secretory vesicles including the micronemes, and a pre-conoidal actin nucleation center (Daher et al., 2010; Dos Santos Pacheco et al., 2022; Martinez et al., 2023). Together, these attributes instruct a unidirectional “apex first” movement of the mature tachyzoite when it egresses out from the host cell. Therefore, there is no need for the usual macroscopic symmetry breaking step to steer the direction of locomotion. Moreover, the curved tachyzoite follows prominently or exclusively a helical path in 2D and 3D conditions, respectively (Leung et al., 2014), which reduces the significant cell-to-cell variability in motile movements commonly observed for most populations.

To start helical gliding on a flat surface, the tachyzoite must adhere to the latter through its base. It then progresses about one body length with a 180° rotation along the major axis, while lifting the apex, and then completes a 360° rotation before the next move forward (Håkansson et al., 1999; Pavlou et al., 2020) (**Fig. 1a**). The long-lasting prevalent mechanistic gliding model is centered on a series of stationary MyosinA (MyoA) motors aligned between the plasma membrane and the inner membrane complex (IMC), itself anchored to 22 cortical spiraled microtubules (cMTs). In this position, the myosin motor units are poised to pull short actin filaments in a fixed rearward orientation, defined by the CMTs’ cytoskeleton, and along the parasite length (**Extended Fig. 1b**). The contribution of cMTs as a template for the F-actin flow was though recently questioned by Hueschen et al. *via* tracking actin filaments in tachyzoites and modeling the emergence of a F-actin apicobasal flow through F-actin self-organization (Hueschen et al., 2022). Force is thought to be transmitted to the ECM across molecular anchoring platforms that are primarily shaped by ECM ligands interacting with the parasite transmembrane adhesin receptors called MICs. MICs are exclusively released at the apical tip of the parasite through regulated exocytosis of micronemes (Ayoade & Chandranesan, 2023; Brown et al., 2016; Chan et al., 2023). MIC2 is acknowledged as the mechano-transducer MIC prototype because dysregulation of its expression drastically alters the tachyzoite helical gliding skills in 2D and 3D conditions (Gras et al., 2017; Huynh & Carruthers, 2006; Shen et al., 2014; Stadler et al., 2022). Upon exocytosis, MIC2 displays an ectodomain featured by an integrin-like A/I-domain and an M domain encompassing six thrombospondin-like repeats. The model supports that a molecular bridge provides the transient mechanical link between F-actin and the adhesins required for force transmission (Jacot et al., 2016; Kumar et al., 2023). Along with the retrograde translocation of the MIC2-F-actin complex, the MIC2 ectodomain is enzymatically cleaved through a multistep process (Carruthers et al., 2000; Zhou et al., 2004), which prevents excessive amounts of bonds between MIC2 at the parasite surface and extracellular ligands at the ECM. These cleavages also permit the timely shedding of the ECM-bound MIC2 molecules from the parasite surface, a strict condition for forward displacement.

**Figure 1.**
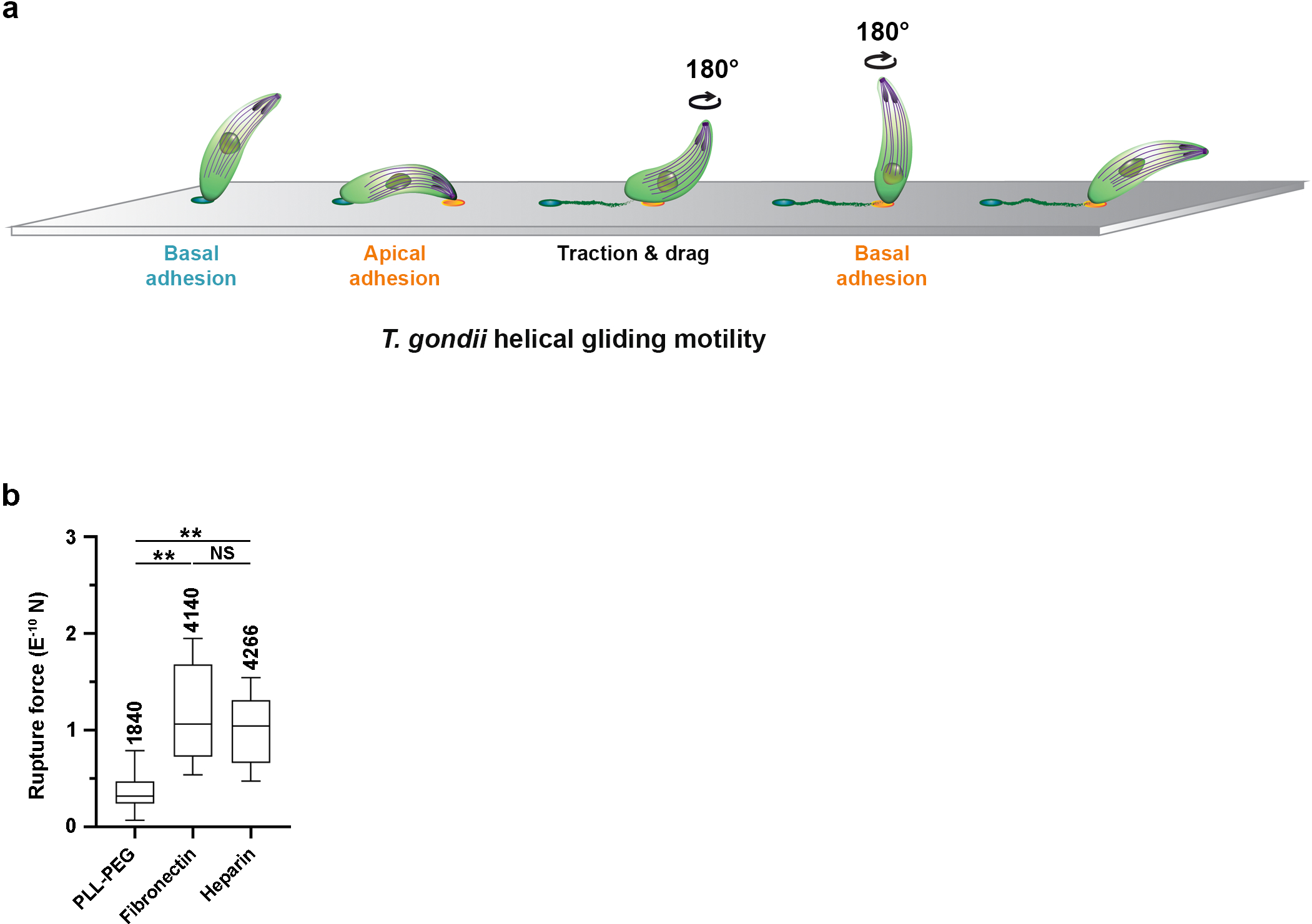
Helical gliding and adhesion forces. **(a)** Schematic of *T gondii* tachyzoite undergoing helical gliding on a substrate. The parasite attaches by the basal pole (blue spot) and builds with its apical pole an anchoring site with the substrate (orange spot) on which it applies traction. Traction drives sliding of the parasite over this site, which proceeds in a helical path with the lift up of the parasite apex while the rear region is drag­ ged forward onto the substrate. This sliding is associated with the release of material shed behind the motile parasite as “trail” (green). Once the basal pole of the parasite has reached the anchoring site, the parasite continues to rotate and repeats the same helical gliding sequence. **(b)** Rupture-force distribution of tachy­ zoites on the PLL-PEG, fibronectin and heparin functionalized surface. Whiskers are maximum and mini­ mum, and boxes indicate second and third quartiles framing the median. Medians for PLL-PEG, fibronectin and heparin are 31.7 pN (2761 cells), 106 pN (13148 cells) and 104 pN (4471 cells), respectively (4 to 5 in­ dependent assays, unpaired t test).

Recently, using reflection interference contrast microscopy (RICM), we have demonstrated that once adhering to the substrate through the basal pole, the tachyzoite involves its apex in a privileged contact site that remains stationary on a flat surface while the tachyzoite progresses over it in a twisting forward motion. This longer-lived contact site serves as an anchor point across which the tachyzoite transmits a MyoA-based traction force that drags the cell body forward (Pavlou et al., 2020). In a related scheme, when gliding within Matrigel or an elastic fibrin-based hydrogel, the tachyzoite must build an adhesive ring-shaped platform that critically supports the transmission of inward-directed forces generated by the parasite to squeeze through, as retrieved from a challenging 3D TFM analysis (Stadler et al., 2022). Therefore, identifying the architecture and composition of these pivotal adhesion platforms is essential to understand how force powers helical gliding.

To this end, we have combined micropatterning, flow force and live or super resolution imaging with tunable surface chemistry, and analyzed the motile capacities of tachyzoites under restrictive adhesion spacing. We have also built well-defined, highly specific and adaptable model surfaces using poly(l-lysine)-g-poly(ethylene glycol) (PLL-PEG) coatings coupled to streptavidin (SAv)/biotin chemistry to tune the nature and density of ligands and to study their effects on parasite motility. We complemented our study with a quartz crystal microbalance with dissipation monitoring (QCM-D) and viscoelastic modeling for biophysical characterization of the tunable surfaces, and identified the ligand species required for gliding. To further challenge the gliding model, we interrogated the relationships between retrograde surface flow, adhesin secretion, and adhesion. Collectively, this work offers a combinatorial framework that permits to experimentally decode the interplay of *Toxoplasma* forces, adhesion and mechanics, and to refine the Apicomplexa gliding mechanistic model.

## RESULTS

### *T. gondii* tachyzoites display similar adhesion forces on heparin and fibronectin functionalized microfluidic chips

Using high-speed RICM, we previously mapped at the sub-micrometer scale the substrate site where the motile parasite must first engage apically to generate a traction force to propel itself forward (Pavlou et al., 2020). However, it remained unclear whether the ECM ligand species encountered by the parasite during this contact can modulate force generation and hence motility. Indeed, no major phenotypic differences in the parasite performance when settled on glass surfaces coated with either proteins (i.e., fibronectin) or complex polysaccharides such as glycosaminoglycans (GAGs) have been reported (Carruthers et al., 2000; Gras et al., 2017; Håkansson et al., 1999; Pavlou et al., 2020). To characterize the overall tachyzoite adhesiveness on biomaterial surfaces, we performed cell-substrate adhesion force rupture assays on large samples of newly egressed parasites deposited on either the high molecular weight plasma fibronectin (*M*w from ∼ 400 to 550 kg/mol) or the negatively charged low molecular weight heparin (*M*w ∼18 kg/mol) immobilized in a microfluidic chip (see Methods). We included in the assay the synthetic poly(l-lysine)-g-poly(ethylene glycol) (PLL-PEG), a hydrophilic copolymer which prevents tachyzoite adhesion (Pavlou et al., 2020). Using brightfield live-cell imaging (3.3 frames/s) of parasites and monitoring their detachment under a controlled flow gradient rate (from a 0.05 to 50 µL/min), it was possible to extrapolate the tachyzoite stalling forces for each surface coating (see Methods). The determined stalling force values, which represent a predominant fraction of resting parasites, were not statistically different between the fibronectin and heparin surfaces (p=0.56, unpaired t test) since a 50% loss of parasites was observed when flow forces reached 106 ± 35 pN and 104 ± 21 pN, respectively. In the case of non-adhesive PLL-PEG, 50% parasites detached at more than 3 times lower stalling forces (32 ± 17 pN) (p< 0.01, unpaired t test) (**Fig. 1b**). While these adhesion rupture assays documented in real time a similar tachyzoite adhesiveness to protein and polysaccharide surfaces at the population level, they did not provide information on whether these individuals can similarly achieve helical gliding on either of these surfaces.

### Polar adhesions to micropatterned substrates are sufficient to support optimal helical gliding of MIC2-competent parasites

To analyze the adhesion requirements at the tachyzoite cell level, and given the necessity of an apical anchoring site for initiating helical gliding (Pavlou et al., 2020), we next interrogated whether this spatially defined site would be sufficient for sustained productive adhesion (i.e., to ensure proper force transmission). Addressing this point allowed revisiting the contribution of the full body adhesiveness during gliding. To this end, we used deep UV printing (Azioune et al., 2010) and designed repetitive rectangle patterns coated with pro-adhesive fluorescent material, fibronectin or heparin, each separated by increasing widths of repulsive PLL-PEG lanes. The length of pro-adhesive rectangles was 20 µm to restrict the chance of helical gliding within the area while still long enough to allow two helical sequences, therefore serving as a good control for the active status of the parasites. The width of these pro-adhesive micropatterns was 2 µm as it is close to the resolution limit of our patterning method and significantly smaller than the parasite length. Indeed, we determined the length distribution of > 600 parasites labeled with the lipophilic membrane dye PKH26 by live imaging to adjust the anti-adhesive PLL-PEG spacing (**Extended Fig. 2a**). Given the 6.7 ± 0.7 µm average parasite length, we designed anti-adhesive PLL-PEG lines of either 4, 6, 8 or 10 µm width (**Extended Fig. 2b**). The 15 min-video sequences of tachyzoites settled on these micropatterns as well as on non-micropatterned fibronectin, heparin and PLL-PEG coatings were analyzed with respect to the distinct motile behaviors among the active tachyzoite fraction **(Fig. 2 and Extended Fig. 2)**.

**Figure 2.**
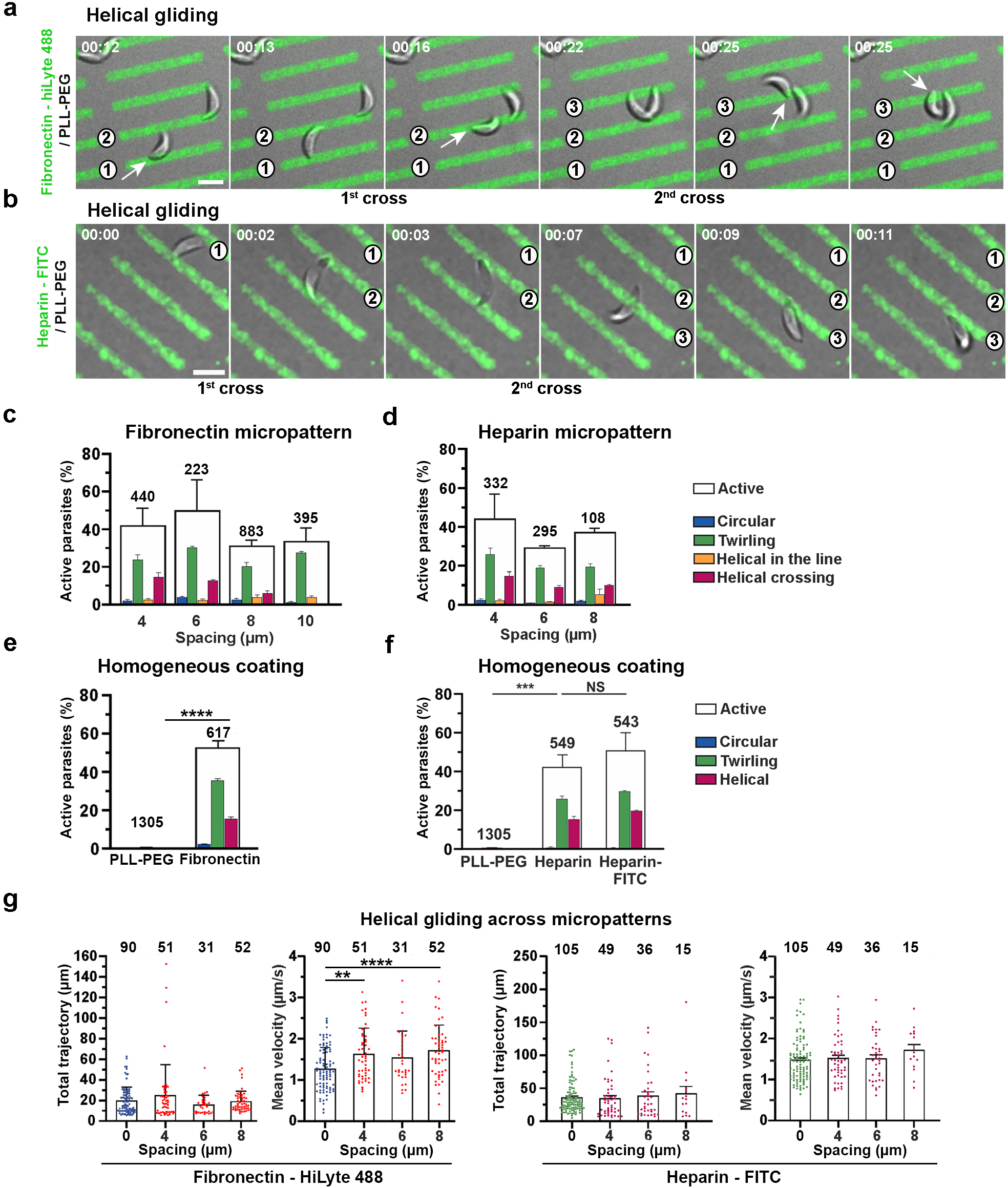
Tachyzoite motile behaviors on geometrically constraint micropatterns. **(a-b)** Bright field-fluorescent overlay images showing tachyzoites that perform helical gliding across rectangular micropatterns (20 µm length x 2 µm width) coated with **(a)** fibronectin-HiLyte488 or (b) heparin-FITC and spaced by 4 µm width lines of anti-adhesive PLL-PEG. Two helical crosses are shown on each surface. The successive pro-adhe­ sive patterns are indicated by numbers. Time is in min:s, scale bars are 5 µm, see also Supplementary movies 1 and 2 for entire time series. **(c-d)** Percentages of active tachyzoites on **(c)** fibronectin or **(d)** hepa­ rin micropatterns. The active tachyzoites are subcategorized according to the three types of movement (i.e., circular, twirling and helical), n = 108 to 883 parasites from at least 3 experiments). **(e-f)** Percentage of active tachyzoites on homogeneous **(e)** PLL-PEG and fibronectin coatings(**** p<0.0001, unpaired t Stu­ dent’s test), **(f)** PLL-PEG and heparin coatings(**** p<0.0001, unpaired t Student’s test). **(g)** Total displa­ cement and mean velocity of tachyzoites performing helical gliding on fibronectin and heparin homoge­ nous coatings (no PLL-PEG = 0 µm spacing) and micropatterned surfaces of (4, 6, 8 µm PLL-PEG spa­ cing). On fibronectin patterns, crossing helical gliding does not differ from gliding on a fully coated co­ verslip in total displacement (NS, p=0.1169, ANOVA followed by Dunnet’s multiple comparisons) whereas the speed of crossing helical gliding is faster (4 µm: ** p<0.01, 8 µm: **** p<0.0001, both ANOVA fol­ lowed by Dunnet’s multiple comparisons). On heparin, tachyzoites display similar total displacement and mean velocity while performing crossing helical gliding on homogeneous or patterned surfaces (n=15-105 parasites from at least 3 independent essays, NS, one-way ANOVA followed by Dunnet’s multiple compari­ sons). For each graph, the number of parasites analyzed are noted on top of the columns. Statistical analyses are detailed in Materials and Methods section.

Tachyzoites were observed to cross from one adhesion permissive rectangle to the next by engaging apically with either fibronectin or heparin micropatterns while undergoing true helical gliding motility **(Fig. 2a-b and Supplementary movies 1-2)**. Overall, the fraction of active tachyzoites retrieved from the motility assays on micropatterns varied between 35 to 50 %, which was slightly below what was measured for tachyzoites settled on homogeneous fibronectin or heparin surfaces (**Fig. 2c to f**) (p=0.20 for fibronectin micropattern *vs* full coat; p=0.78 for heparin micropattern vs full coat, **Supplementary Table 2**; the difference in p likely comes from the reduced sample size in the heparin micropattern *vs* full coat). In all cases, the predominant fraction of motile tachyzoites (more than 50%) twirled upright around their main axis with the basal pole attached to the substrate, whereas less than ∼ 8% underwent circular trajectory. However, up to 33 ± 13% and 32 ± 5% tachyzoites were able to glide across fibronectin and heparin micropatterns, respectively (**Fig. 2c-d**) while the frequency of crossing events decreased with the PEG spacing to reach zero for the 10 µm width (**Fig. 2c**). To detail the behavior of parasites on the PLL-PEG/fibronectin (or PLL-PEG/heparin) micropatterns, we next compared the helical gliding performances of tachyzoites on the homogenous and patterned surfaces. While no significant difference was detected in total displacement (p=0.12), the mean velocity of parasites was found somewhat faster on PLL-PEG/fibronectin patterns (p<0.0001), an effect that was however not reproduced on heparin (**Fig. 2g**).

In addition to the cell-repellent PLL-PEG copolymer, we also used poly(N-isopropylacrylamide) (PNIPAM) brushes to generate fibronectin rectangular micropatterns separated by PNIPAM lines of 6 and 8 µm width with a submicron resolution. PNIPAM can be grafted to surfaces covalently and at high density, providing coatings of strong protein and cell repellency at 37°C (Mandal et al., 2012), while allowing a strong contrast in RICM (Varma et al., 2016). Similar to PLL-PEG, PNIPAM prevented tachyzoite adhesion since the parasites only attached to the adjacent fibronectin areas. Aside from twirling on fibronectin, the tachyzoites could cross over the PNIPAM lanes from one fibronectin area to the next (**Fig. 3a and Supplementary movie 3)**. Video-rate (20 Hz) 4D joint reconstruction of the micropattern-parasite interface brought definitive evidence that the apical contact made by the tachyzoite with the fibronectin micropattern during the crossing event allows sufficient surface anchorage to withstand the forward displacement traction, while the rest of the parasite body remains above the substrate. In contrast, when the parasite apex got close to the PNIPAM area, proper anchorage could no longer form, thus preventing gliding, and the parasite lifted up, twisted on its base and tried again (**Fig. 3b and Supplementary movie 4**).

**Figure 3.**
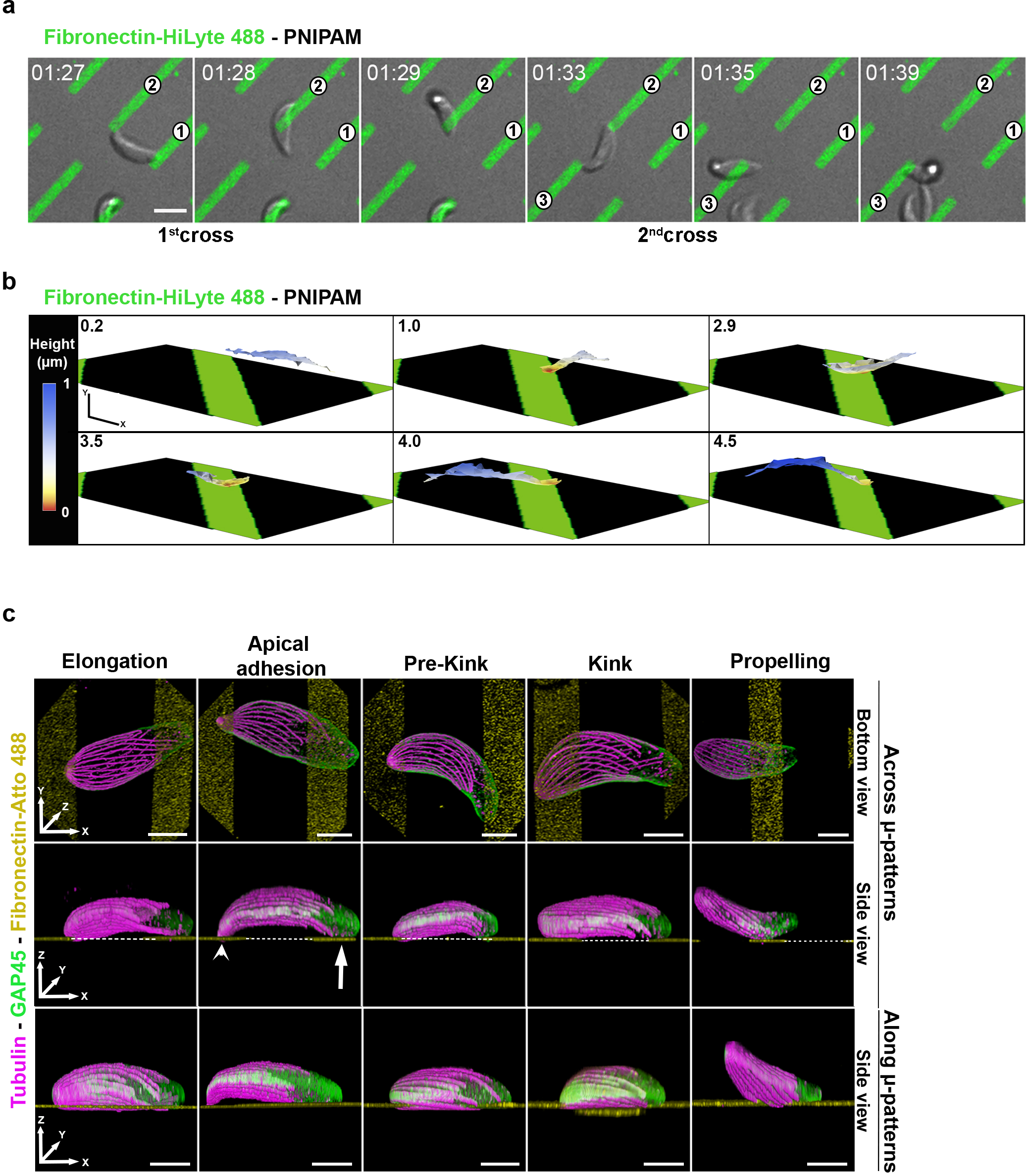
Spatiotemporal mapping of the parasite-surface contacts during helical gliding across pro-adhesive micropatterns. **(a)** Representative bright field-fluorescent overlay images showing a tachyzoite that per­ forms two helical gliding sequences across rectangle (2×20 µm) micropattems coated with fibronectin-Hi­ Lyte488 and spaced by 6 µm width PNIPAM brushes. Time is in min:s, scale bar is 5 µm, see also Supple­ mental movie 3 for entire time series. **(b)** 3D reconstruction of RICM images of fibronectin-PNIPAM mi­ cropattems (green) and of the bottom surface of the parasite body (red to blue, color-coded as a function of the distance to the coverslip) during one helical gliding sequence and a subsequent failed attachment onto PNIPAM. Adhesive 2 µm rectangles are spaced by 6 µm of PNIPAM. 6 representative images were ac­ quired at 20 Hz over 10 s. Time indicated on each image is ins. X scale bar= 2 µm, Y scale bar= **1** µm. See also Supplementary movie 4 for entire time series. **(c)** Confocal fluorescent images of tachyzoites ex­ panded by U-ExM while undergoing helical gliding by crossing over anti-adhesive 4 µm wide PLL-PEG lanes that separate fibronectin-Atto488-coated rectangles (2 µm, yellow areas in top and central rows) or by moving along the pro-adhesive rectangle (yellow areas in bottom row). Each frame recapitulates a different step of the helical sequence and shows tachyzoites immunostained for the submembranous GAP45 glideo­ some component (green) and for the cortical microtubules (magenta). White arrowhead and arrow mark the apical and basal contacts, respectively at the moment when the tachyzoite makes a polar contact with the neighboring fibronectin rectangle. The top row presents a bottom view while the central and bottom rows present side views of the 3D image reconstruction; scale bars are 2 µm, see also Supplementary movies 5 and 6 for entire volume reconstruction.

Since the MIC2 protein has been characterized as the crucial adhesin to productively engage with ECM ligands during gliding (Huynh & Carruthers, 2006; Gras et al., 2017), we monitored the behavior of tachyzoites genetically lacking the *mic2* gene (Δ*mic2*) (Gras et al., 2017) on the fibronectin micropatterns. As expected, tachyzoites lacking the MIC2 protein showed much lower activity compared to the MIC2 positive parasites on both fibronectin (p=0.0016) and heparin (p=0.0001), with a similar ∼ 5-times loss of motile capacities on both surfaces (p=0.12), while keeping twirling as a major motile behavior (**Extended Fig. 2c)**. In line with these results, only very few MIC2 deficient tachyzoites were able to cross over PLL-PEG area of 4 μm width to reach either a fibronectin-or heparin-coated micropatterns, with only one crossing event (1 parasite out of 357) being observed on the 6 μm heparin micropattern and none on the 8 μm fibronectin and heparin ones (**Extended Fig. 2d**). These results position MIC2 as a primary adhesin candidate for the assembly of the force-productive platform driven by the initial contact with the pro-adhesive surface.

### Ultrastructure-expansion microscopy highlights differences in the parasite-surface contact features on micropatterned and uniform surfaces

To study how the tachyzoite undergo helical gliding on a homogenous surface, we have recently coupled RICM with ultrastructure expansion microscopy (U-ExM) and proposed that spiral cMTs bend locally and transiently under the actomyosin contractile force when the tachyzoite builds apical contact with the surface and starts gliding (Pavlou et al., 2020). Once the compression is released, the cMTs transiently straighten, while the parasite disengages apically from the substrate and moves one length of its body forward. To get *in situ* nanoscale insights on parasite-substrate contacts during helical gliding, we have applied U-ExM (Gambarotto et al., 2021) to fibronectin-Atto^488^ micropatterns. Fibronectin-Atto^488^ conjugate allowed delineating micropatterns on expanded samples while their initial fixed dimension (2, 4, 6, 8 μm) provided direct control for the correct *xy* isotropic specimen enlargement. We could “freeze” tachyzoites as they moved either across PLL-PEG towards a neighboring fibronectin patch or along a fibronectin area. Post-expansion labeling of the parasite contours and cortical microtubules (cMTs) allowed visualizing with ∼60 nm resolution the contact surface between the parasite and the micropattern as well as the curvature of the spiral cMTs.

Using this approach, we have reconstructed the helical gliding multistep sequence based on the RICM profile previously defined in Pavlou et al., 2020 (**Fig. 3c**). Our present analysis confirmed that tachyzoites made only polar contacts with the two successive fibronectin areas by extending their body, while keeping their natural curvature with their concave side away from the PEG area. Some crossing tachyzoites bent apically on the fibronectin area (i.e., likely corresponding to the kink step, Pavlou et al., 2020) for which we observed a tight contact between the parasite apex and the beneath micropattern with the apically deformed cMT network (**Fig. 3c, 4^th^ frame)**. Additionally, the apically uplifted parasites with straight microtubules were caught at the end of their sliding on the established anchorage on the micropattern (**Fig. 3c, 5^th^ frame)**. Interestingly, parasites moving across micropatterns (**Fig. 3c, central panel, Supplementary movie 5**) were found to flatten less on the substrate than those gliding along fibronectin areas (**Fig. 3c, bottom panel, Supplementary movie 6**). The pronounced flattening of the latter matches the RICM and TFM data obtained on homogenous coatings, which identified the progressive increase in the contact area between the parasite and the substrate as the tachyzoite body drags during its traction (Pavlou et al., 2020). In the case of micropatterned surfaces, the reduced contact area between the substrate adhesive molecules and the parasite should result in weaker friction, which may be one of the reasons for the increased speed of tachyzoites moving across micropatterns (**Fig. 2g**). Collectively, these results confirm the polar adhesion requirements for the parasite twist along the spiral cMTs during helical progression. They also suggest that a reduced frictional interface imposed by the pro-adhesive micropattern geometry could account for the observed increase in the parasite gliding velocity.

### Designing specific and tunable model surfaces with physico-chemical properties characterized by QCM-D

We next explored the molecular characteristics of the substrate sufficient for the productive apical anchoring of the parasite. Pro-motile conditions for *T. gondii* tachyzoites include a pro-adhesive substrate as illustrated with micropatterns, but also proper trigger(s) for the secretion of MIC adhesins (Brown et al., 2016; Bullen et al., 2016). Indeed, most parasite motility assays have been carried out in presence of serum albumin in the medium or/and pre-adsorbed on glass surfaces. In addition, both lipid-bound and lipid-free serum albumin molecules have been characterized as highly potent and selective secretagogues of soluble and transmembrane MIC proteins, acting in synergy with the activation of the protein kinase G pathway (Brown et al., 2016). With the goal of identifying the ECM-derived ligands that would engage with the parasite-exposed adhesins and support force transmission during helical motility, it was crucial to design an assay in which we could control the nature of molecular species exposed to the parasite, while preventing adsorption of serum albumin or other serum components to the surface. To monitor the buildup of such tunable model surfaces as well as to characterize the specificity of parasite/surface interactions on the previously studied coatings, we used quartz-crystal microbalance with dissipation monitoring (QCM-D). QCM-D is an acoustic surface-sensitive technique, which allows highly sensitive real-time monitoring of the adsorption kinetics, binding stability and specificity through measuring the resonance frequency (*Δf*, reflecting the mass uptake) and dissipation (*ΔD*, related to the film softness) shifts of the piezoelectric sensor (**Fig. 4a**).

**Figure 4.**
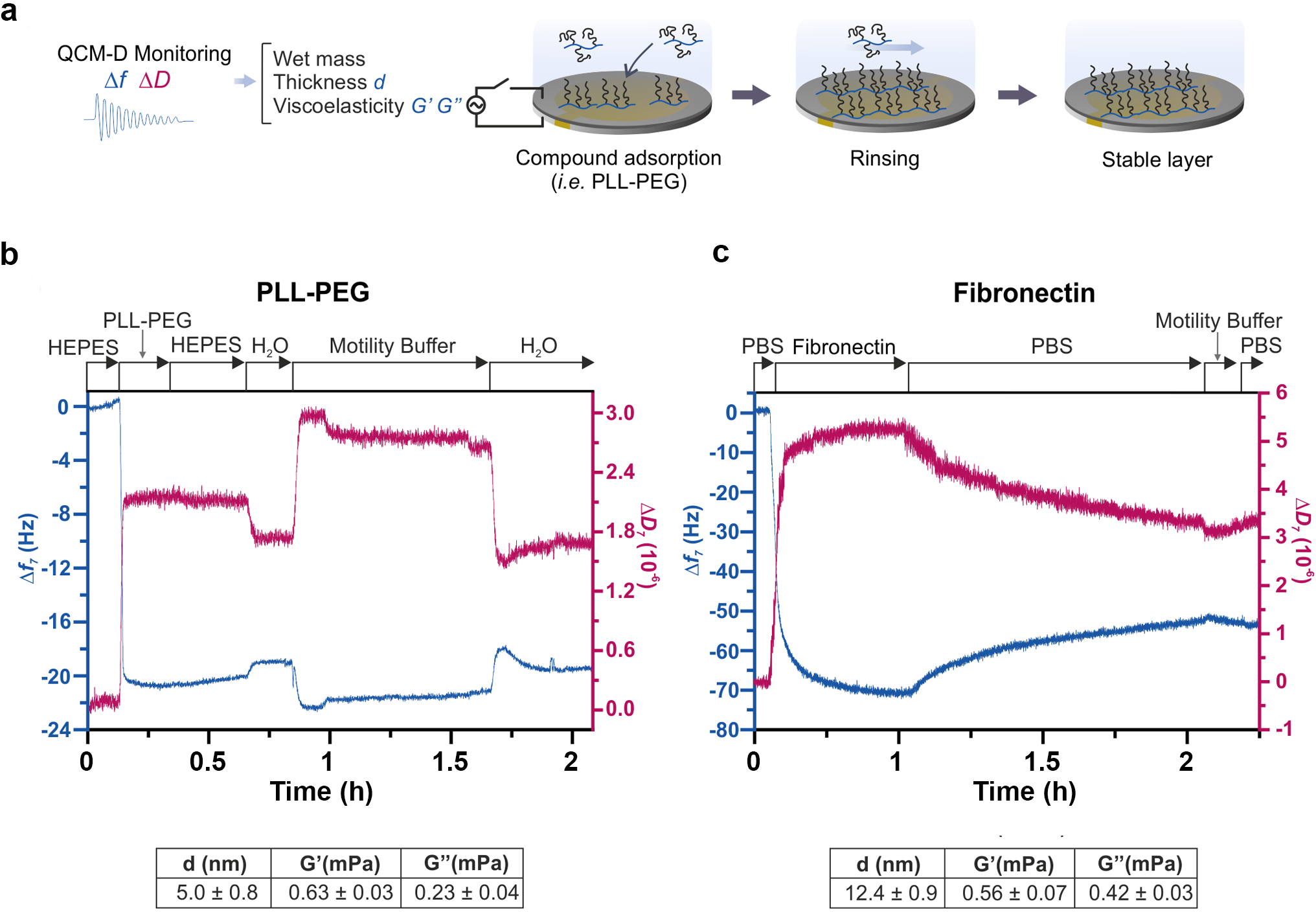
QCM-D characterization of molecular coatings used in micropatterns. **(a)** Schematic of QCM-D measurements. The silica-coated quartz sensor oscillates at a resonance frequency, allowing to monitor shifts in frequency (llf) and dissipation (llD) upon adsorption on its surface. Measurements are perfor­ med in solution, and the stability of molecules attachment is verified upon rinsing with water or working buffer. The solutions in contact with the surfaces at different moments of time are indicated on the top of QCM-D profiles. **(b-c)** Characteristic llf and llD shifts (7th overtone) obtained for PLL-PEG (b) and fibronectin **(c)** binding to silica. In both cases, the components of the tachyzoite motility buffer do not attach to the formed layers as evidenced by the absence of llf and llD shifts upon their injection and rin­ sing. Thickness **(d),** storage modulus (G’) and loss modulus **(G″)** values determined using viscoelastic modeling are presented as mean± SEM (n=4 independent assays for PLL-PEG and n=5 independent assays for fibronectin) below the QCM-D graphs.

QCM-D analysis documented strong adsorption of the motility buffer (MB) components on the widely used bare **(Extended Fig. 3a)** and PLL-coated silica **(Extended Fig. 3b**) surfaces. In contrast, no binding from the components of the tachyzoite MB (e.g., FCS, glucose) was observed on the PLL-PEG and fibronectin coatings formed on the silica surface (**Fig. 4b-c**). In addition, QCM-D results confirmed that PLL-PEG binding reached saturation very fast (< 5 min) and resulted in a stable and soft (Δ*D* > 0) coating (**Fig. 4b**). QCM-D monitoring of fibronectin binding to silica indicated rather slow binding (saturation in ∼1 h), resulted in a soft (Δ*D* > 0) coating, a major fraction of which was stable in studied media (**Fig. 4c**). Furthermore, to compare physico-chemical parameters of the repellent PLL-PEG and adhesive fibronectin layers, we determined their thicknesses (*d*) and viscoelastic properties (i.e., storage modulus *G’* and loss modulus *G’’*) by viscoelastic modeling of QCM-D data. The results showed that the two layers had similar storage moduli while the thickness and loss modulus of the fibronectin layer (12.4 ± 0.9 nm and 0.42 ± 0.03 mPa) were 2.5- and 1.8-times higher than that of PLL-PEG (5.0 ± 0.8 nm and 0.23 ± 0.04 mPa, inset tables in **Fig. 4b-c**). The concomitant changes in these parameters suggest that not only the chemical nature, but also the thickness and softness of the molecular interface may contribute to the parasite motile capacities. Of note, the observed *Δf* and *ΔD* profiles as well as the viscoelastic modeling performed were consistent with the ones previously reported (Hemmersam et al., 2008; Perry et al., 2009; Saftics et al., 2018).

Capitalizing on the PLL-PEG surface results, we have developed a tunable model system based on streptavidin (SAv)/biotin chemistry, enabling to compare the specific effects of the nature and density of the solution-facing molecules on tachyzoite adhesiveness and motility. To this end, we first attached PLL-PEG-biotin (PPL-PEG-b) to silica surfaces, followed by the binding of SAv:biotin-X (SAv:b-X) complexes, where X represents different functional groups or molecules (**Fig. 5a**). Thanks to the tetravalent nature of SAv, one can tune SAv:b-X ratio in solution, which allows varying b-X density on the surface upon SAv:b-X attachment. To compare different functional groups, we selected mono-biotinylated versions of PEG (3-5 kg/mol) with a terminal hydroxyl (-OH), amine (-NH_2_) or carboxylic group (-COOH). Besides, we studied mono-biotinylated versions of molecules with reported effects on tachyzoite motility, such as human serum albumin (HSA) and heparan sulfate (HS) (Huynh & Carruthers, 2006). The efficiency of surface functionalization and the specificity of SAv/biotin interactions were verified by QCM-D. The adsorption of PLL-PEG-b to the silica surface was fast (∼5 min), resulting in a stable and soft coating (Δ*D* > 0) (**Fig. 5b**), similarly to PLL-PEG (**Fig. 4b**). QCM-D showed successful binding of SAv to the PLL-PEG-b layer with a saturation achieved within ∼15 min, resulting in stable and rigid coating (Δ*D* ∼ 0). A higher Δ*f* = −39.5 ± 0.5 Hz was found for SAv binding to PLL-PEG-b compared to well-studied biotinylated self-assembled monolayers and supported-lipid bilayers with typical shifts of −25-30 Hz (Dubacheva et al., 2017), confirming formation of highly dense SAv monolayers on flexible biotinylated PLL-PEG coatings. No SAv binding was observed on PLL-PEG lacking biotin (**Extended Fig. 4a**) as well as none of the studied b-X molecules interacted with PLL-PEG-b in the absence of SAv (see example **in Extended Fig. 4b**), confirming the specificity of SAv/biotin interactions in our system. When SAv:b-X complexes were bound to PLL-PEG-b, higher Δ*f* and Δ*D* shifts were observed (**Fig. 5b triangles, Fig. 5c and Extended Fig. 4c,d**) as compared to SAv alone (**Fig. 5b circles**), proving the presence of specific b-X constructs on our surfaces. As expected, the magnitude of Δ*f* and Δ*D* shifts as well as the time needed for their saturation increased with the size of b-X constructs (**Fig. 5b-c**). Remarkably, higher QCM-D shifts were detected when increasing SAv:b-X ratio from 1:1 to 1:2 (e.g., > 2-times higher Δ*D* in the case of 1:2 as compared to 1:1 SAv:b-PEG-OH and SAv:b-HS), confirming that stoichiometry of SAv:b-X complexes can be controlled in solution and, as a result, the density of b-X ligands can be controlled on the surface. Importantly, the components of the buffer used in our gliding assays (glucose, FCS, salts) did not interact with any of the PLL-PEG-b/SAv:b-X layers built. Indeed, negligible Δ*f* and Δ*D* shifts were detected upon exposure of PLL-PEG-b/SAv:b-X coatings to the motility medium (**Fig. 5b’-c’, Extended Fig. 5 c’-d’-e**), implying that all potential effects on parasite motility must be due to the presence of specific b-X ligands. Finally, viscoelastic modeling of QCM-D data allowed characterizing the thickness and viscoelasticity of the PLL-PEG-b/SAv:b-X layers. Similar thickness, storage and loss modulus were obtained for PLL-PEG-b/SAv:b-X coatings, in which X represents PEG-OH, PEG-NH_2_ and PEG-COOH of same size (3-5 kg/mol) and at the same stoichiometry (1:2), proving the robustness of the viscoelastic fitting protocol **(Fig. 5d**). Together, these data highlight several advantages of the developed model surfaces over earlier tachyzoite adhesion assays, including control of the specificity of parasite/surface interactions, the ability to tune the nature and density of surface ligands, and to quantify the thickness and viscoelastic properties of the produced coatings.

**Figure 5.**
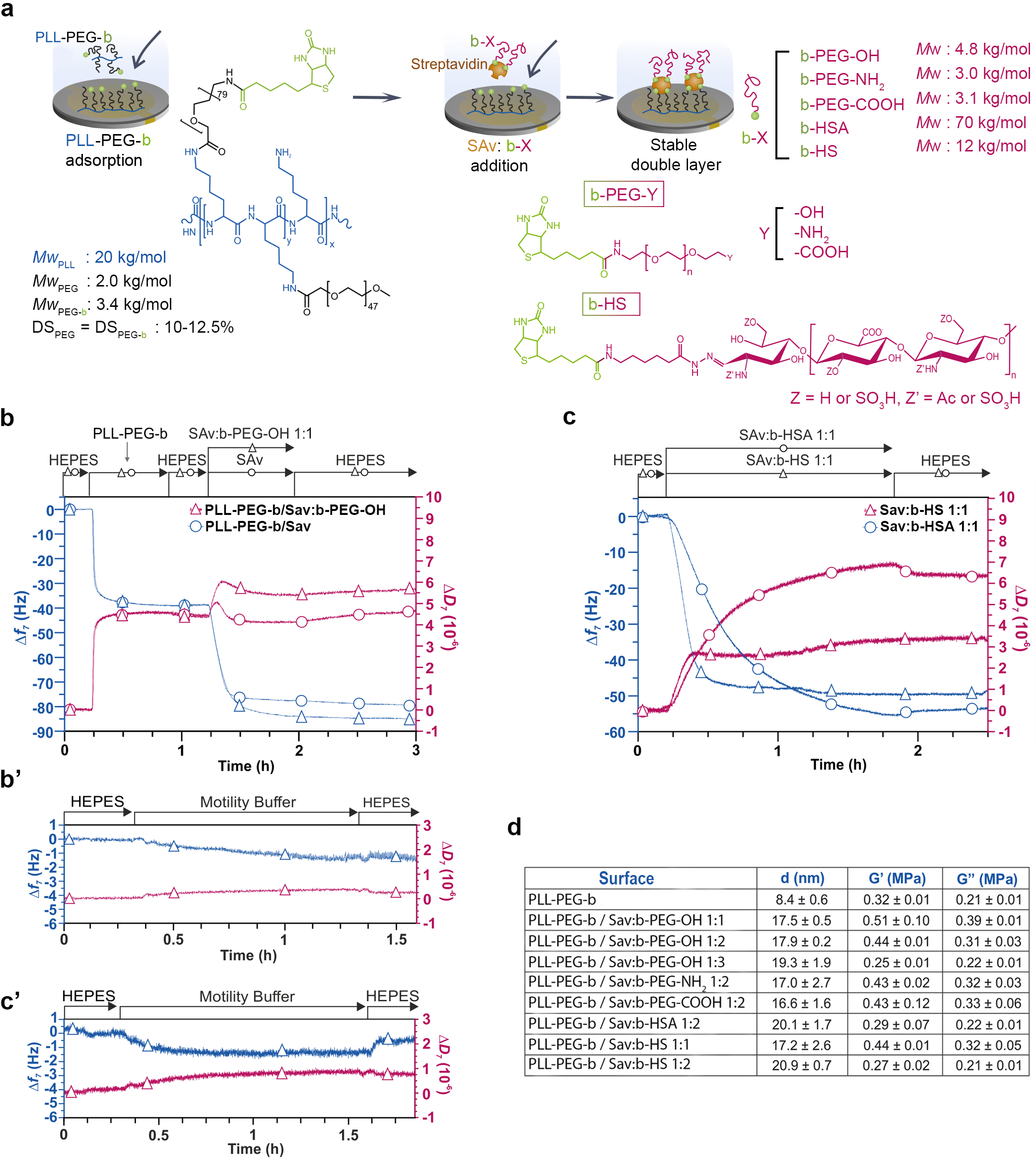
Buildup of tunable model surfaces using PLL-PEG and SAv/biotin chemistry monitored by QCM-D. **(a)** Main steps of surface functionalization, allowing tuning the terminal molecule while using the same sur­ face chemistry, are shown together with chemical structures and molecular weights of the studied ligands. First, PLL-PEG-biotin (PLL-PEG-b) is adsorbed on the silica-coated sensor, followed by the injection of streptavidin/biotin-X (SAv:b-X) complex, being X the tunable ligand. The solutions in contact with the sur­ faces at different moments of time are indicated on the top of QCM-D profiles. **(b)** Characteristic llf and *ii* D shifts (7th overtone) recorded upon PLL-PEG-b binding to silica, followed by the attachment of SAv (cir­ cles) or SAv:b-PEG-OH **(1:1)** complex (triangles). **(c)** Characteristic QCM-D profiles recorded upon the at­ tachment of SAv:b-HS **(1:1)** (triangles) and SAv:b-HSA **(1:1)** (circles) complexes to the PLL-PEG-b layer preformed on silica. **(b’, c’)** The components of the tachyzoite motility buffer do not show strong attach­ ment to the PLL-PEG-b/SAv:b-PEG-OH **(1:1) (b’)** and PLL-PEG-b/SAv:b-HS **(c’)** coatings; with same re­ sults being obtained for the PLL-PEG-b/SAv:b-HSA system (see Extended Fig. 4e). **(d)** Thickness **(d),** sto­ rage modulus *(G’)* and loss modulus *(G’’)* determined using viscoelastic modeling. Values represent mean± SEM from n = 2-17 independent QCM-D experiments.

### Glycosaminoglycans are sufficient ligands to promote productive interaction with tachyzoite adhesins for helical gliding

To identify the minimal molecular ligand requirements for the tachyzoite to engage with the substrate in a productive contact, we first confirmed that tachyzoites do not adhere, hence do not show any gain of motile capacity on PLL-PEG-b and PLL-PEG-b/SAv surfaces (i.e., lacking the specific ligands) when compared to the anti-adhesive PLL-PEG (p=0.13; **Supplementary Table 2**) (**Fig. 6a**). Next, we tested the tachyzoite motile response to the presence of serum albumin that is commonly used in parasite motility assays and was shown to influence the secretion of MIC proteins (Brown et al., 2016). Using biotinylated human serum albumin (b-HSA, 70 kg/mol) as a ligand, we observed a restoration in the tachyzoite active phenotype when compared with the PLL-

**Figure 6.**
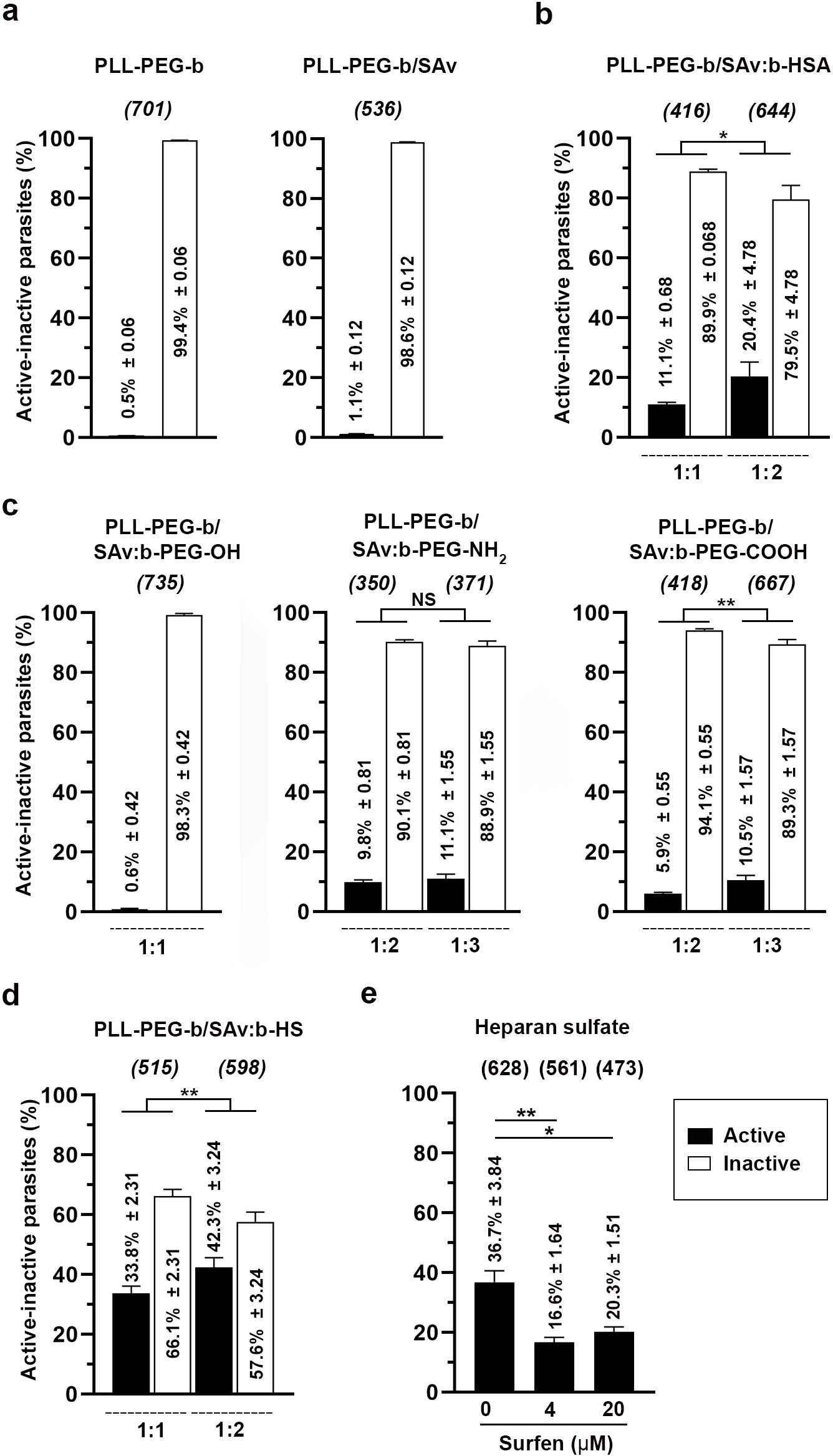
Tachyzoite activity on tunable surface coatings characterized by live imaging. **(a)** Tachyzoites are not active on PLL-PEG-b and PLL-PEG-b/SAv coatings lacking specific ligands. **(b)** The presence of b-HSA ligands increases the activity(**** p<0.0001, Fisher’s exact test). Doubling the density of exposed moie­ ties additionally promotes activity(*, p=0.0306, Fisher’s exact test). **(c)** For the b-PEG ligands with tunable terminal groups, -OH does not allow to gain motile capacities, while -COOH and -NH2 do with a signifi­ cant increase in the activity (**** p<0.0001, Fisher’s exact test). Increasing -COOH surface density in­ creases tachyzoite motility even further (**, p=0.0091, Fisher’s exact test). (**d)** A significant increase of pa­ rasite activity is observed in the presence of b-HS ligands(**** p<0.0001, Fisher’s exact test). Doubling the b-HS surface density increases the activity even further(**, p=0.0055, Fisher’s exact test). The activi­ ties measured in (b-d) are compared to the non-adhesive PLL-PEG coating (Fig. 2e). See Fig. 5a for chemi­ cal structures and molecular weights of the studied biotinylated ligands. **(e)** Charge neutralization of the di­ rectly attached to the surface heparan sulfate coating with surfen decreases the tachyzoite activity (**, p=0.0030 and*, p=0.0176 comparing 0µM to 4µM and 20µM, respectively) with similar effects being ob­ served in the studied concentration range (4-20 µM, NS, p=0.1804, unpaired t test). For all graphs, bars re­ present mean± SEM from at least 3 independent experiments, the number of parasites analyzed are noted between parentheses. Statistical analyses are detailed in the Materials and Methods section and Supplemen­ tary Table 2.

PEG conditions (p<0.0001). Interestingly, an increase in the HSA surface density noticeably boosted the tachyzoite motile response (p=0.03) (**Fig. 6b**). These results confirm that the presence of serum albumin on surfaces affects tachyzoite behavior, stressing the importance of suppressing non-specific interactions with the tachyzoite MB components (**Fig. 5b’-c’ and Extended Fig. 5c’-d’-e**) for the identification of surface ligands promoting tachyzoite motility.

We next probed the effect of the surface charge on the parasite motile capacities. When the tachyzoites were exposed to the polar PEG-OH ligands, the permissiveness of the substrate to gliding was not statistically different from that of PLL-PEG (p=0.66). In contrast, the presence of charged -NH_3_^+^ or -COO^-^ terminal groups on PEG chains (pH of the MB is of 7.2) resulted in a modest fraction of active tachyzoites (< 10%), which nevertheless was statistically different from the anti-adhesive PLL-PEG (p<0.0001) (**Fig. 6c**). To further explore the effect of charge, we tested the tachyzoite motile response to the presence of the highly negatively charged heparan sulfate, which was shown to be a relevant ligand for *T. gondii* motility and invasion (Carruthers et al., 2000). Indeed, using biotinylated heparan sulfate (b-HS) as a ligand in our model system induced the highest motile response, which was further increased with the b-HS surface density (statistical difference between 1:1 and 1:2 ratio, p=0.005) (**Fig. 6d**). Furthermore, we analyzed the effect on tachyzoite motility of neutralizing negative charges on the substrates using the small molecular antagonist surfen (*bis*-2-methyl-4-amino-quinolyl-6-carbamide) known to electrostatically bind heparin, heparan sulfate and other GAGs. Surfen binding was shown to neutralize the ability of heparins to activate antithrombin but also to alter a range of biological processes mediated by the heparan sulfate (Schuksz et al., 2008). Exposure of silica surfaces coated with heparan sulfate to surfen significantly reduced the gliding ability of tachyzoites (p=0.0016, one-way ANOVA) (**Fig. 6e**). These results confirm the important contribution of electrostatic interactions to promote a productive focal adhesion for motility on surfaces coated with negatively charged GAGs. In addition, viscoelastic modeling of QCM-D data demonstrated that while the thickness and viscoelastic properties of some PLL-PEG-b/SAv:b-X coatings were very similar (for instance, see the results obtained for SAv:b-PEG-OH (1:1), SAv:b-PEG-NH_2_ (1:2), SAv:b-PEG-COOH (1:2) and SAv:b-HS (1:1), **Fig. 5d**), the motile responses of tachyzoites varied significantly on them (p<0.0001, Fisher’s exact test) (**Fig. 6b-c-d**). This confirms the predominant contribution of the biochemical nature of the ligand over that the mechanical properties, at least in the stiffness range of our experimental settings. In summary, our results identify the negative charge of ECM-exposed ligands as an essential condition to engage in a productive interaction with the parasite adhesins to enable gliding.

### Sensing extracellular condition rather than parasite adhesion to the surface is required to trigger surface protein flow

In the classical 2D gliding model, attachment of the parasite to the surface is considered as a trigger for the apically directed exocytosis of microneme vesicles (Carruthers et al., 1999). This vesicle fusion event leads to the release and incorporation of the transmembrane MIC adhesins at the plasma membrane, a process thought to be tightly coupled with the activation of an actomyosin driven apicobasal membrane flow, itself required for motility (Carruthers et al., 1999). Regulatory mechanisms of microneme protein secretion include the apical trafficking of micronemes (Chan et al., 2023; Lentini et al., 2019; Leung et al., 2017) and signaling cascades involving calcium or/and cGMP second messengers upstream of the exocytosis event (Brown et al., 2016; Bullen et al., 2016; Li et al., 2022). In order to validate the proposed contribution of substrate adhesion as a trigger for apicobasal flow (Carruthers et al., 1999), we performed high-speed imaging (200 Hz) to compare in physiological thermic conditions (37°C) the behavior of 0.2μm-diameter microbeads captured (i) by active parasites on fibronectin (i.e., extruding their conoid and subsequently gliding) or (ii) by those settled on PLL-PEG, thus unable to adhere and glide.

The tachyzoites settled on fibronectin were seen to respond to the MB by extending the conoid and capturing the microbeads. As shown in **Fig. 7a and Supplementary movie 7**, a single microbead that came in contact with the conoid translocated readily to the basal pole, around which it rotated further just before the parasite lifted up while undergoing twisting forward motion. The microbead was eventually released behind the parasite when the latter underwent helical propelling. This immediate apicobasal motion of the bound bead implies that the directionality of the parasite’s sub-membranous actomyosin system, which powers the membrane protein flow, must have been functional at the time the conoid and body extended and prior to the parasite motion. Consistent with this finding, we established that the apicobasal translocation of microbeads from the tip to the basal end also occurred when parasites were exposed to MB and settled on PLL-PEG. Yet, because the parasite could not pull on the substrate, the translocated microbead stayed attached to the base even after 26 seconds (**Fig. 7b and Supplementary movie 8**). Despite selecting only active parasites that extruded their conoid and further engaged in typical helical movement on fibronectin, we found that the average speed of the microbead varied significantly (3.90 ± 1.3 µm/s, n=14). In the case of the parasites “floating” on top of the anti-adhesive PLL-PEG layer, the average speed of microbead translocation was found to be somewhat lower (1.99 ± 0.36 µm/s, n=5). Overall, the measured velocities of microbead apico-basal motion are slightly higher than the parasite’s gliding speed (1-2 µm/s) (Pavlou et al., 2020), and largely exceed those previously found using the OTs setting. Of note, we did not observe the random motion reported in Stadler et al. (2017) to precede the polarized movement of a microbead manipulated with optical tweezers (OTs). This difference likely lies in the OTs approach that was used on nonactive parasites adhering lengthwise to the PLL-coated surface (i.e., conoid retracted, immotile) at 24°C.

**Figure 7.**
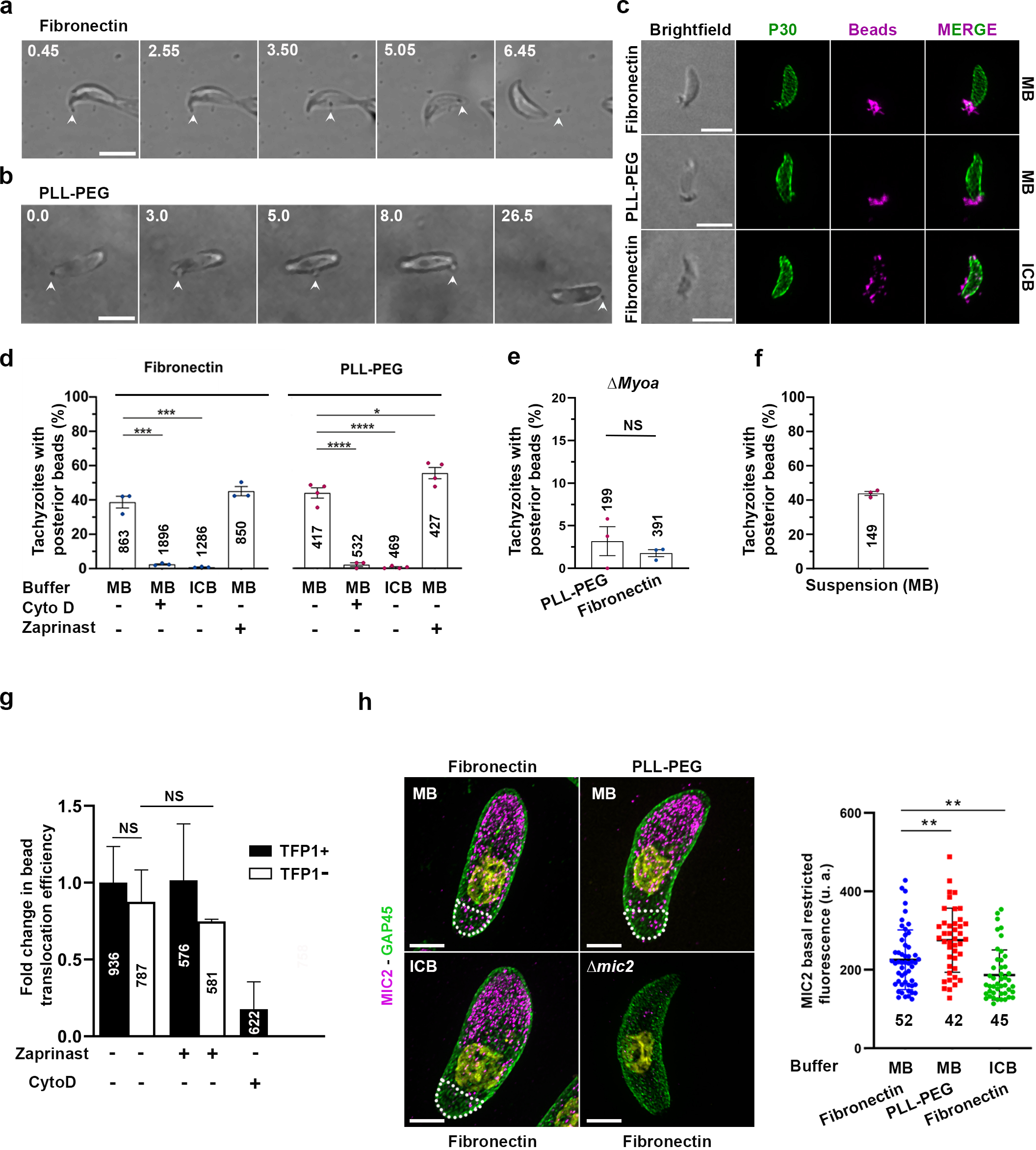
Surface protein flow characterized by monitoring translocation of surface-bound microbeads and re-localization of MIC2 adhesins. **(a, b)** Representative bright field images showing a tachyzoite whose apical tip engages in a contact with a microbead, which is then translocated to the basal pole and **(a)** is further released while the tachyzoite moves forward on fibronectin or **(b)** remains associated when the parasite is unable to glide on PLL-PEG. The white arrowheads mark the microbeads. Time is in min:s, scale bars are 5 µm, see also Supplementary movies 7 and 8 for entire time series. **(c)** Representative confocal images showing bright field views (1st column) and the maximal intensity projections of tachyzoites (2nd, 3rd, 4th columns) with bound microbeads (magenta) and immunostained for the surface glycoprotein P30 (green). Clustering of beads at the posterior pole indicates an efficient surface surface protein flow when exposed to MB (two first rows) and settled on fibronectin (top row) or PLL-PEG (median row). In contrast, the micro-beads dispersed over the whole cell surface support the absence of surface protein flow when tachyzoites are exposed to ICB and settled on fibronectin (last row). Scale bars are 5 µm. **(d, e, f)** Percentages of tachyzoites with posterior microbeads indicate: **(d)** high efficiency of surface protein flow for cells settled on either fibronectin or PLL-PEG and exposed to MB or MB supplemented with zaprinast, with very low efficiency detected upon disruption of actin dynamics (cytochalasin D/cytoD) or when exposed to ICB, **(e)** surface protein flow is almost abolished for **Δ**myoA tachyzoites settled on either fibronectin or PLL-PEG and exposed to MB, **(f)** efficient surface protein flow occurs when parasites are maintained in suspension in MB. Numbers of scored parasites indicated in the graphs are from at least 3 independent assays. Statistical analyses are detailed in the Materials and Methods section and Supplementary Table 2. **(g)** Percentages of tachyzoites expressing TFP1 (TFP1+) or conditionally depleted in TFP1 (TFP1-) with posterior microbeads when settled on fibronectin and exposed to MB or ICB, and treated or not with CytoD or Zaprinast as indicated below the graph. Numbers of scored parasites indicated in the graph columns are from at least 3 independent assays. Statistical analyses are detailed in the Materials and Methods section and Supplementary Table 2. **(h)** Maximal projection images of MIC2-expressing tachyzoites exposed to MB or ICB and settled on fibronectin or PLL-PEG as well as of **Δ**mic2 tachyzoites settled on fibronectin and exposed to MB. U-ExM-expanded samples immunostained for GAP45 (green), MIC2 (magenta), DNA (yellow) display MIC2-positive patches at the posterior region (white dotted line), which suggest microneme exocytosis and MIC2 apicobasal translocation in response to MB independently of adhesion. Scale bars are 2 µm. Quantification of MIC2 signal pixel intensity at the posterior region is shown on the right graph together with numbers of analyzed parasites. Statistical analyses are detailed in the Materials and Methods section and Supplementary Table 2.

To further quantify the efficiency of the apicobasal microbead translocation process, we enumerated the fraction of tachyzoites whose entire surface was initially covered with fluorescent microbeads in intracellular mimicking buffer (ICB), and for which a cluster of microbeads was detected at the posterior pole after 5 min of incubation in MB buffer (**Fig. 7c**). Comparing proadhesive (fibronectin) and anti-fouling (PLL-PEG) surfaces, the fraction of tachyzoites with translocated microbeads was found comparable (**Fig. 7d**) and similarly dependent on a competent actomyosin system, as assessed using either the actin polymerization inhibitor cytochalasin D (p<0.0001) or tachyzoites genetically deficient in the MyoA motor (**Fig. 7d-e**). In the same line, in the presence of MB, the tachyzoites mechanically maintained in suspension translocated beads with a similar efficiency as those settled on surfaces (**Fig. 7f**). Interestingly, the tachyzoites responded positively to zaprinast (100 µM) added to the MB to overstimulate microneme secretion (C. R. Collins et al., 2013). Indeed, an increased number of tachyzoites with posteriorly clustered microbeads was observed on PLL-PEG (p=0.0212) and to a less extent on fibronectin (p=0.3842). Besides, almost no tachyzoites were able to posteriorly cap microbeads when exposed to the intracellular mimicking conditions on either pro- or anti-adhesive surfaces (p<0.0001) (**Fig. 7d)**. Collectively, these results demonstrate that surface “protein” flow, assessed *via* the translocation of 0.2 μm bound microbeads, occurs through the same actomyosin mechanisms under proadhesive and anti-adhesive conditions. They also highlight that the extracellular buffer conditions which include fetal serum, millimolar amounts of calcium and glucose are sufficient to set up the actomyosin-driven flow machinery, therefore priming the tachyzoite for its subsequent ability to glide.

### Microneme exocytosis and adhesin release at the cell surface are not required for the apicobasal translocation of surface bound-microbeads

The ability of the parasite to translocate microbeads posteriorly in absence of contact with the surface, and the slight increase of translocation efficiency in conditions that boost microneme secretion (i.e., Zaprinast) (**Fig. 7d**), prompted us to revisit the relationships between microneme exocytosis and surface flow. To this end, we took advantage of the parasite line genetically engineered to express the tetracycline-induced system for conditional depletion of the gene encoding the microneme associated transporter family protein 1 (TFP1). In the absence of TFP1 expression, tachyzoites are fully impaired for microneme exocytosis and unable to glide or invade cells (Hammoudi et al., 2018). The efficiency of TFP1 depletion and its effect on the parasite motility were assessed 42h post tetracycline (ATc) addition by *in situ* immunostaining of both parental and silenced parasites and by scoring their motile and cell invasive capacities (**Extended Fig. 5a-b**). We compared the amounts of TFP1-expressing and TFP1-depleted parasites that translocated microbeads when exposed to MB and settled on fibronectin. Remarkably, we found that the TF1-depleted tachyzoites were able to translocate microbeads with similar efficiency than the parental TFP1-expressing tachyzoites (p=0.70) (**Fig. 7g**). These data rule out the view that the microbead retrograde flow reflects the process of microneme exocytosis and adhesin release at the cell surface.

Since microneme exocytosis was found to be unnecessary for the actin-dependent surface protein flow process (**Fig. 7g**), we next analyzed the integration of the MIC2 adhesin in this flow, while comparing pro- and anti-adhesive substrates (**Extended Fig. 2c)**. Using U-ExM, we resolved MIC2 patchy signals and confirmed their highest density at the apical pole of tachyzoites. We interpreted patches as individual micronemes, but could not reliably discriminate the intracellular micronemes from those that have undergone exocytosis and have likely incorporated into the plasma membrane hence exposing adhesins. To quantify the efficiency of MIC2 rearward trafficking, we analyzed the fluorescent signals detected at the posterior region of tachyzoites, mostly devoid of intracellular micronemes using an antibody against the cytoplasmic C-terminal region of MIC2 (see Methods). Given that tachyzoites barely translocate microbeads in ICB **(Fig. 7d)**, we expected to detect fewer MIC2 patches at the basal region of tachyzoites exposed to ICB.

Indeed, a significant reduction of MIC2 posterior fluorescent signal characterized the tachyzoites incubated in ICB as compared to MB buffer (p=0.0029) when both were settled on fibronectin (**Fig. 7h**). Importantly, the number of MIC2 posterior patches was even slightly higher in the case of parasites exposed to MB but settled on PLL-PEG compared to those settled on fibronectin (p = 0.0081) (**Fig. 7h, graph**). We attribute this difference to the fact that the parasites are unable to glide on PLL-PEG and thus cannot lose the MIC2-displaying material in the trails left behind, a scheme that recalls the longer persistence of the microbead at the basal pole when tachyzoites were settled on PLL-PEG (**Supplementary movies 7 and 8**).

Collectively, these results argue for a microneme exocytosis independent- and actomyosin-dependent mechanism of surface protein apicobasal flow, which further incorporates adhesins upon microneme exocytosis. Therefore, how the membrane pools are delivered apically and further recycled posteriorly remains an open question. It has recently been reported that dense granules are discharged subapically at the annuli site, including during the extracellular life (Chelaghma et al., 2023), a process down-regulated by zaprinast (Katris et al., 2019). We therefore tested the effect of this compound on the capacity of TF1-lacking tachyzoites (i.e. impaired in microneme exocytosis) to translocate microbeads when exposed to MB. We found that TFP1-lacking parasites exhibited similar efficiency of the apicobasal protein surface flow regardless of the use of zaprinast when assessing posterior translocation of microbeads (p=0.8) (**Fig. 7g)**. These observations indicate that dense granules are unlikely the source of apical membrane required to drive the microneme-independent but actin-dependent protein surface flow.

### Tachyzoites respond to an excess of microneme secretion by a persistent twirling behavior

Zaprinast, which boosts microneme secretion, was reported to induce a larger fraction of MIC2-expressing tachyzoites to engage in motility on fibronectin (Lourido et al., 2012). In contrast, it was shown that tachyzoites cannot tract themselves forward when impaired in their ability to disengage their basal pole from the substrate, either as a result of MIC2 posterior accumulation (Buguliskis et al., 2010) or artificial immobilization setting (Pavlou et al., 2020). Therefore, we interrogated whether the tachyzoites could sense and respond to the detrimental excess of surface exposed adhesins in order to regain gliding capacities. Using live imaging, we determined that the primary behavioral response to zaprinast was a robust twirling behavior. Indeed, in less than 30 s post addition of zaprinast to MB, about 71% (n =283, 2 independent assays) of tachyzoites engaged in twirling, while a sub-fraction of those underwent helical or circular gliding in the 5 min-period of recording on fibronectin substrates (**Supplementary movie 9**). As expected, the tachyzoites settled on PLL-PEG or PNIPAM did not exhibit gliding, yet about a third of them started to display a persistent twirling motion in response to zaprinast present in MB (PLL-PEG: n= 99/301, 3 independent assays; PNIPAM: n= 58/161, one assay) (**Supplementary movie 10**). In this twirling subset on PLL-PEG, most tachyzoites were still jointed posteriorly through the residual body and leaned against each other to apply torques, but a few could also succeed in twirling alone. These observations suggest that the accumulation of proteins and lipids at the parasite rear, caused by the drug-induced microneme secretion, has increased the chances of insertion of adhesin-rich biomaterial within the repellent coatings, and consequently provided enough anchoring for sustaining twirling. These data also uncover a close relationship between the microneme exocytosis-driven surface flow process and the twirling behavior, which might indicate that the parasite senses and tunes the adhesion strength at the posterior pole to improve their gliding capacities.

## DISCUSSION

When gliding across surfaces, the *T. gondii* tachyzoite progresses with a 360° helical forward movement in an iterative manner, with each period lasting a few seconds. Each of these periods includes an obligatory molecular engagement of the parasite apex with the substrate to sustain the generation of an actomyosin traction force that couples with adhesion disassembly at the cell rear (Pavlou et al., 2020). However, quantitative information on the adhesive requirements for successful gliding is missing. In this study, ∼100 pN adhesive forces of tachyzoites settled on either fibronectin or heparin and only in part being engaged in motility due to the increasing flow speed, support the view that *T. gondii* has evolved a strategy to restrict body stickiness to the substrate. Yet, to optimize the chance to rapidly access and penetrate the host cell, they must timely generate sufficient adhesion strength to resist the fluid dynamics that characterizes each tissue at homeostasis, and to undergo gliding (Polacheck et al., 2014).

To investigate the minimal requirements for productive adhesion, we chose to design rectangle micropatterns that alternate non-adhesive and pro-adhesive molecular coatings, as a way to geometrically control the contact permissive area proposed to the tachyzoite. Real-time monitoring of the tachyzoite behavior on these micropatterns allowed demonstrating that if attached through the basal pole, the cell needs only one new apical contact site with the substrate to then slip on it through additional transient (possibly nonspecific) interactions that keep it on track. Both 3D and 4D nanoscale mapping of the parasite-surface contact during gliding using U-ExM and high speed RICM identified the limited slippage interface when gliding across micropatterns, which likely decreases the overall frictional interaction zone and favors speed increase when the cell body is dragged forward. In the same line, decreased adhesion of the sporozoite morphotype from *Plasmodium berghei*, phylogenetically related to *T. gondii*, has been reported to correlate with increased speed (Münter et al., 2009). Of note, lower adhesion conditions have been experimentally imposed to metazoan cells using substrates engineered with sparse molecular ligands for the major family of surface-exposed adhesins (integrins). Alone or in combination with confinement between flat substrates, these lower adhesion settings led to increased crawling speed of dermal fibroblasts, which even switched to amoeboid motility under combined conditions (Liu et al., 2015).

Aside from the substrate characteristics, the distribution, life time and activity of the adhesins at the cell surface are key parameters to fine tune parasite adhesiveness. Since the MIC adhesins are exclusively released at the apical tip of the parasite and further processed along the cell surface *via* the apicobasal translocation, both events shape an apicobasal MIC gradient and the “leading edge” of the cell. In 2D and 3D conditions, MIC2 has been identified as the major adhesin that controls gliding (Gras et al., 2017; Stadler et al., 2022), and our micropattern motility assays unambiguously support the MIC2 mechano-sensor function. In addition, using microbeads and MIC2 surface protein flow markers coupled with high-speed live imaging and complementing our experiments with assays on mutant parasites unable to secrete adhesins at their surface, we have uncovered that the actomyosin-dependent cortical flow occurs independently of cell attachment to the substrate and of adhesin secretion, and is likely independent of dense granule secretion, but requires extracellular conditions. In other words, it is the sensing of the extracellular conditions that primes the parasite for future motility by organizing the F-actin system for apicobasal movement of surface proteins. Of note, virtually nothing is known on the complex lipid-protein composition and distribution of the tachyzoite plasma membrane. In particular, the dynamics of each molecular repertoire and their interplay are far from being understood. For instance, an intriguing actin-independent retrograde membrane flow has been detected by monitoring the movement of 0.4 μm microbeads attached to the *T. gondii* surface, which was attributed to the balance of apical exocytosis and basal endocytosis (Gras et al., 2019). Similarly, the impact of microneme exocytosis on the membrane flow dynamics remains elusive. Besides, molecular dynamic simulation and quasi-elastic neutron scattering experiments have been applied to simplified membranes, and the obtained data support a scenario where transmembrane proteins slow down lipid dynamics (Ebersberger et al., 2020). Moreover, the anchorage of transmembrane proteins to the cortical actin cytoskeleton promotes the confinement of the former in nano/microdomains which have been reported as sources of membrane’s resistance to flow in many cell types (Shi et al., 2018). In the case of *T. gondii*, we found that the speed of the translocating microbeads is about two times faster than the speed of parasites. This suggests that the interaction between the substrate ligands and the secreted transmembrane adhesins impacts the F-actin flow while promoting the assembly of a transient focal adhesion platform and the buildup of adhesive force. Such scheme is supported by studies on the *Plasmodium* sporozoite which, interestingly, also exhibits about twice faster microbead translocation when compared to gliding speeds when using OTs (Quadt et al., 2016).

In the context of a minimal adhesion scenario, we also propose that the twirling response observed upon zaprinast addition could reflect the sensing of an excess of membrane material capped posteriorly, due to the overloaded processes of MIC endocytic recycling and/or enzymatic cleavage. In turn, the tachyzoites engage mechanically their posterior poles with the surface (including the residual body or a paired tachyzoite) in a specific torque to lower the interfacial adhesive strength at their basal pole and thereby improve chances for further gliding activity. Indeed, a complete loss of gliding capacity was reported when the parasite was unable to disengage from the posterior pole (Pavlou et al., 2020), while preventing MIC2 enzymatic processing post-secretion was found to impair gliding and favor twirling (Buguliskis et al., 2010). Accordingly, our observations argue that trails are by-products of the microneme secretion process due to incomplete recycling and pile-up of membranous material posteriorly, which forms and extends when the parasite pulls on the anchoring adhesion site and slips over it.

The next important question is how the MIC2 adhesins engage with their ligands to build the minimal anchor point. Several early structural studies of MIC2 have established that the MIC2 A domain binds heparin provided that the former is presented as large, multimeric complexes (Harper et al., 2004; Tonkin et al., 2010). In the same line, MIC2 was reported to be expressed as a hetero-hexamer comprised of three MIC2-MIC2-Associated Proteins (MIC2AP) and three MIC2 molecules on the surface of *T. gondii* with the idea that multivalent interaction would enhance ligand-adhesin binding efficiency (Jewett & Sibley, 2004). By combining tunable surface chemistry with QCM-D, we could build highly specific interfacial platforms with well characterized biophysical properties and adjustable molecular settings, including the nature and the density of molecular ligands. We observed that in the absence of molecular adsorption from the medium onto the surface, the presentation of negatively charged ligands like GAGs to the parasite is sufficient for promoting a parasite-substrate productive contact, in which electrostatic interactions are likely one of the driving forces for coupling adhesins to GAG. In particular, the behavior of parasites on heparan sulfate was shown to be density dependent, as increasing the number of exposed moieties resulted in the higher fraction of motile parasites. In addition, we showed that the surfaces with the same viscoelastic parameters but with different ligands (e.g., PEG *vs* heparan sulfate) result in drastically different motility characteristics. This suggests that ligand chemistry may have a superior influence on the capacity of tachyzoite to glide as compared to other interfacial parameters. *In vivo*, heparan sulfates are known to bind to numerous growth factors and ECM proteins, including fibronectin and HS controls fibronectin assembly in fibrils serves as the base for building the ECM network in many tissues (Raitman et al., 2018). In addition, heparan sulfates are abundantly secreted by endothelial cells and circulate as fragments in the blood, in particular during inflammation (Collins & Troeberg, 2018). Therefore, tachyzoites must be continuously exposed to heparan sulfates when navigating within acellular compartments and interacting with the cell glycocalyx during gliding or invasion. The approach we have developed allows dissecting parasite interactions with heparan sulfates as well as the contributions of other proteoglycans or glycoproteins forming ECM, and thus uncover the related ligand-receptor requirements for *T. gondii* host cell invasion. Furthermore, combining our combinatorial toolkit with additional surface-sensitive techniques and an extended range of surface chemistries (e.g., allowing lateral mobility and/or nanostructuration) should allow to address a larger set of questions including quantification of the effects of ligand density, membrane fluidity, and clustering on *T. gondii* adhesion and motility.

## MATERIALS AND METHODS

### Molecules used for surface functionalization

**PLL** (P8920, Sigma Aldrich), **PLL-PEG** (poly-L-lysine-grafted-polyethylene glycol PLL (20 kDa) - g[3.5] - PEG (2 kDa), SuSoS); **PLL-PEG-b** (biotinylated poly-L-lysine-grafted-polyethylene glycol PLL (20 kDa) - g[3.5] - PEG (2 kDa) / PEG (3.4 kDa) - biotin (50% of PEG chains), SuSoS); **fibronectin** (bovine plasma fibronectin, F4759, Sigma Aldrich, ∼ 450 kg/mol); **fibronectin-HiLyte^488^** (FNR02 Cytoskeleton, ∼ 230 kg/mol); **heparin** (H3149-100KU, Sigma Aldrich, Mw ∼18 kg/mol); **heparin-FITC** (H7482 ThermoFisher, ∼ 18 kg/mol); **heparan sulfate** (sodium salt from bovine kidney, H7640, Sigma Aldrich, ∼ 10-14 kg/mol); **SAv** (streptavidin S4762, Sigma Aldrich, ∼60 kg/mol); biotinylated ligands (b-X): **b-PEG-OH** (PEG1207.0100, Iris Biotech, 4847 g/mol), **b-PEG-NH_2_** (PEG1046, Iris Biotech, 2996 g/mol), **b-PEG-COOH** (PEG1053, Iris Biotech, 3145 g/mol), biotinylated human serum albumin **b-HSA** (70 kg/mol, ACROBiosystems HSA-H82E3, Fisher Scientific), biotinylated heparan sulfate **b-HS** (synthesized using hydrazone ligation to 12 kg/mol HS and characterized by QCM-D to determine % of biotinylation as described in Thakar et al. 2014).

### Buffers

**HBSS^++^**: Hanks balanced solution (137.9 mM NaCl, 5.33 mM KCl, 0.34 mM Na_2_HPO_4_, 0.44 mM KH_2_PO_4_, 4.1 mM Na HCO_3_, 1.26 mM CaCl_2_, 0.49 mM MgCl_2_, 0.40 mM MgSO_4_), pH 7.2. Motility buffer **(MB)**: HBSS^++^ buffer supplemented with 1% fetal calf serum (FCS) and 1.6 mM CaCl_2_, pH 7.2. Intracellular mimicking buffer **(ICB)**: 145 mM KCl, 5 mM NaCl, 1 mM MgCl_2_, 15 mM MES, 15 mM HEPES, pH 8.3. **HEPES buffer**: 10 mM HEPES, pH 7.2 or 7.4.

HEPES-NaCl buffer: 10 mM HEPES, 150 mM NaCl, pH 7.4. **Carbonate buffer**: 0.1 M NaHCO_3_/Na_2_CO_3_, pH 8.3. Phosphate buffer saline (**PBS**): dPBS 1X without calcium and magnesium, 14190-144, Gibco, pH 7.4. **MES buffer**: 50 mM MES, pH 6.0. **U-ExM Denaturation buffer**: 50 mM Tris, 200 mM SDS, 100 mM NaCl, pH 9.0. **PHEM fixation buffer**: 60mM PIPES, 25mM HEPES, 10mM EGTA, and 4mM MgSO_4_.

### *T. gondii* tachyzoite maintenance and handling

-Maintenance: *Toxoplasma gondii* strains were propagated by serial passages on *Mycoplasma*-free human foreskin fibroblasts (HFFs, ATCC CCL-171) grown in T25 cm^2^ flask at 37 °C and 5% CO_2_ atmosphere in the complete medium [Dulbecco’s modified Eagle’ medium (DMEM) supplemented with 10% heat-inactivated fetal bovine serum (Invitrogen), HEPES buffer (pH 7.2), 2 mM L-glutamine, 100 U/mL penicillin, and 50 µg/mL streptomycin)]. All reagents were from Gibco - Life Technologies (St Aubin, France). The *T. gondii* RH *KU80* knockout strain (Δ*ku80*) was provided by V. Carruthers (University of Michigan, USA). The RH Δ*ku8*0: DiCre ΔMyoA (Δ*myoA)* and RH Δ*ku80*:diCre/Δ*mic2* (Δ*mic2*) were provided by M. Meissner (MLU Munich University, Germany) (Gras et al., 2017; Huynh & Carruthers, 2009). The RH Δ*ku80* Tati TFP1-3Ty-iKD line (TFP1-iKD) was provided by D. Soldati-Favre (University of Geneva, Switzerland)(Hammoudi et al., 2018).

-Handling: for motility assays, tachyzoites were collected within a few hours of spontaneous egress from the HFF monolayers, which were carefully synchronized for infection (1 to 2 parasite/cell). After slow short centrifugation (200g, 1 min, to remove cell debris) of 1 mL supernatant obtained from the 5 mL culture, the tachyzoite-containing supernatant (i.e., ∼3×10^6^ tachyzoites) was transferred to a 50 mL tube and centrifuged in a large volume of HBSS^++^ (800g, 7 min). The pellet was resuspended in MB for assays. For the membrane flow assays that required intracellular mimicking buffer conditions, the monolayer of *T. gondii*-infected HFF cells in T25 flask were cultured for ∼ 40 h to obtain 32 to 64 tachyzoites loaded intracellular vacuoles but no parasite egress. The culture medium was then replaced by two successive washes in ICB and the monolayer was gently scrapped in 2 mL of ICB. The collected material was passed through a 25G syringe to release the intracellular tachyzoites. The suspension was then diluted in 45 mL of ICB and collected by serial centrifugations as described above, after that the pellet was resuspended in 2 mL of ICB and immediately used in the assays. To silence the expression of TFP1, the Tati TFP1-3Ty-iKD parasites were produced on HFF monolayers (T25cm^2^ flasks) that were exposed immediately after infection and for about 44 to 48 h to 1 μg/mL Anhydrotetracycline (631310, TaKaRa, ATc) whereas control cultures were exposed to vehicle only (100% ethanol).

### Tachyzoite motility and invasion assays

-Motility assays: to minimize cell debris, the parasite suspension prepared as described above was additionally filtered through a hydrophilic membrane of 5 μm pore size (Cyclepore Tracked Etched Membrane, Cytiva, Fisher Scientific). The functionalized 18- or 32 mm diameter glass coverslips (micropatterned or not) were placed inside the microscope chamber and preequilibrated in MB at 37°C and in 5% CO_2_, after that ∼ 1-3×10^5^ tachyzoites were added dropwise onto the 150 or 300 μL of MB covering the coverslips, respectively. Images were captured at 2 to 15 frames/s depending on the microscope and assay. A minimum of 3 video-sequences of 5 min each were taken for each surface/condition, and from 3 independent coverslips and tachyzoite cultures. Image analysis for characterizing tachyzoite activity (active/inactive), type of movement (twirling, circular and helical; helical displacement across or along micropatterns), and tracking displacement as well as velocity of gliding was done using Icy (de Chaumont et al., 2012), with plugins “Manual tracking” and “Motion profiler” to extract the *x-y* positions and time.

-Invasion assays: 12 mm diameter glass coverslips were seeded with HFF to form a monolayer. 1×10^5^ of TFP1 expressing or TFP1-silenced tachyzoites were deposited on a coverslip in complete medium. After synchronized sedimentation (2 min, 300g), the cells were incubated (15 min) at 37 °C and in 5% CO_2_ atmosphere. Extracellular parasites were flushed out with excess of cold PBS before fixation of the HFF samples with 3.2% paraformaldehyde (PFA, 15714, Electron Microscopy Sciences) in PHEM fixation buffer (20 min, RT). Samples were next processed for immunostaining to score invasiveness by quantifying intracellular parasites in the two samples as described in the “Lipid membrane staining and immunostaining of fixed samples” section.

### *T. gondii* tachyzoite adhesion assay using laminar flow shear force

The microfluidic channel (50×5×0.1mm) was deposited on the glass slide (0.1mm μ-slide, Ibidi). The channel was successively cleaned with 2% hydrochloric acid (10 min) and 10% sulfuric acid (5 min). The glass surface was activated using 33% hydrogen peroxide (10 min) and then washed with water and equilibrated with PBS (10 min). Next, the glass surface was exposed to 50 μL of PLL-PEG (100 μg/mL), fibronectin (10 μg/mL) or heparin-FITC (10 μg/mL) in PBS and incubated 3h at room temperature. Unabsorbed reagents were washed away with 200 μL of PBS. The microfluidic chamber was warmed at 37°C prior to aspirate the suspension of tachyzoites prepared as described above in MB (∼ 1×10^5^ cells in 50 μL, 37°C) into the functionalized channel using a syringe. Tachyzoites were incubated 10 min to allow sedimentation and then exposed to a laminar shear flow with a linear gradient rate from 0 to 39.13 mL/min (6.52 10^-7^ m^3^/s) in 90 s, providing a dragging force from 0 to 8.41 10^-10^ N on a spheroid of 1.8 μm-diameter (Juzans et al., 2020). The flow was created using a 60 mL syringe (Omnifix Braun) controlled by a computer-monitored pump (KDS 410). Up to 2500 tachyzoites were imaged with a camera (ORCA, Hamamatsu) in the field of view of an inverted transmission microscope (Observer D1, Zeiss) with an objective 10x/Na 0.5 on a heated stage (37.2 digital Zeiss). Images were acquired (3 images/s), after that cells were counted every 10 images using ImageJ to extrapolate the dragging force *vs* flow rate. The rupture forces were calculated as described previously (Kamsma et al., 2018).

### PLL-PEG and PNIPAM micropatterning

Glass coverslips were cleaned in 0.5 M HCl (overnight, 65°C) and extensively washed in water and ethanol prior to be used. Micropatterning with PLL-PEG was done as follows. The coverslips were exposed to photochemical oxidation by UV/ozone (λ=230 nm, UVO Cleaner, Jelight) for 5 min to improve polymer adhesion, and the activated surface was immediately coated with 0.1 g/L PLL-PEG in HEPES buffer (pH 7.4) (30 min, room temperature). The PEGylated coverslips were then placed onto the chrome side of a UV/ozone-activated patterned quartz photomask (5 min) (UXM1, Toppan Photomasks inc., Texas USA) using a few μL of water as a capillary contact. The PLL-PEG was irradiated through the micropatterns drawn on the photomask by deep UV light for 5 min at a fixed distance of 5 cm from the UV light which allowed submicrometer spatial resolution. The micropatterned coverslips were recovered and the burned regions were coated with pro-adhesive molecules as described in the next section. In the case of poly(N-isopropylacrylamide) (PNIPAM), the micropatterning protocol included the following steps. (i) Glass coverslips were cleaned with piranha mixture (3 vol. H_2_SO_4_:1 vol. H_2_O_2_) for 1h, rinsed with water and blow dried. (ii) A monolayer of aminopropyltriethoxysilane (APTES) was grafted on the coverslips by immersing the substrates in a 0.1% solution of APTES in toluene for 15 min. (iii) The APTES monolayer was further derivatized in a 2.5% solution of 2-bromo-2-methylpropionylbromide in dichloromethane for 5 min, yielding a surface-immobilized coating acting as an initiator layer for radical polymerization. (iv) The initiator layer was selectively de-activated by irradiation of the coverslips with deep-UV light (UVO Cleaner, Jelight) through the photomask for 1 min. (v) A PNIPAM brush (dry thickness ∼20 nm, swollen thickness ∼40-50 nm), corresponding to grafted chains of *M*w ≈ 35 kg/mol, was finally grown from the patterned initiator layer by Atom Transfer Radical Polymerization with in situ catalyst regeneration as described in (Varma et al., 2016).

### Functionalization of micropatterns with fibronectin, heparin and heparan sulfate

The obtained PLL-PEG and PNIPAM micropatterned coverslips were incubated (active side down) on a drop of fibronectin (15 μg/mL) mixed with fibronectin-HiLyte^488^ (10 μg/mL) in carbonate buffer or heparin-FITC (10 μg/mL) in PBS for 30 min at room temperature. The homogenous coatings were obtained by incubating the illuminated side of UV/ozone activated glass coverslips with fibronectin (25 μg/mL in carbonate buffer), heparin (10 μg/mL in PBS), or heparan sulfate (5 μg/mL in 10 mM HEPES pH 7.4) for 40 min at room temperature. The functionalized coverslips were gently rinsed with carbonate buffer (or PBS or HEPES pH 7.4) and equilibrated in MB before being used for the motility assays. In the case of heparan sulfate neutralization, heparan sulfate-functionalized coverslips were exposed for 45 min to 4 or 20 μM surfen (S6951, Sigma Aldrich)(Schuksz et al., 2008) in HEPES buffer (pH 7.4), rinsed in the same buffer then in MB and immediately used for videorecording.

### Dynamic and static microscopy on micropatterned and homogeneous functionalized surfaces or cell monolayers

-Time-lapse videomicroscopy was performed in Chamlide chambers (Live Cell Instruments, Seoul, Korea) accommodating 18 or 35 mm-diameter glass coverslips, installed on an Eclipse Ti inverted confocal microscope (Nikon France Instruments, Champigny sur Marne France) at 37°C and 5% CO_2_ environment (chamber and controllers are from LCI, Seoul, Korea). The microscope was equipped with a piezo element and PrismBSI express camera (Photometrics), a live SR module of super resolution (Gataca), and a CSU X1 spinning disk (Yokogawa). Videos and images were taken using either a 60X Plan-neofluor (1.46 NA, Zeiss) or a 40X Pan Fluor (1.30 NA, Nikon) oil immersion objectives using the Metamorph software (Universal Imaging Corporation, Roper Scientific, Lisses, France). Epifluorescence images were acquired using a 63X Plan-Apotome oil immersion objective (1.46 NA, Zeiss) in an AxioImager M2 fluorescence upright microscope (Zeiss). Image analysis was performed with Metamorph, Icy Software from the raw *x,y,z,t* stacks (https://icy.bioimageanalysis.org, de Chaumont et al., 2012) and Fiji for tridimensional representation (Schmid et al., 2010). Photocompositions were made using Fiji and Photoshop software unless specified in following sections. All images are representative of independent replicates.

-For the microbead/ flow fast monitoring, high-resolution brightfield imaging was performed on an Olympus IX71 inverted microscope (Olympus, Japan) equipped with a 60x, 1.35NA oil objective (Olympus) and an ORCA Flash4 v2 camera (Hamamatsu, Japan) in a thermoregulated chamber. Images were taken at a 200 Hz and further processed by resampling one every 10 images prior to particle tracking analysis.

-For the real time RICM microscopy, the microscope and objective were the same as used for microbead trafficking analysis, but a broadband light (HPLS300, Thorlabs) was used instead of epi-illumination after passing through a multiband dichroic filter (FF01-457/530/628-25, Semrock). Both blue (λ457 nm) and green (λ530 nm) reflection images were detected side-by-side on the camera using a home-built image splitter. Images were recorded at 20 Hz and processed with ImageJ to retrieve the height profile of the lower boundary of the parasite body using custom-written routines based on the Weka Segmentation plugin (Arganda-Carreras et al., 2017). Subsequent tridimensional representation was achieved with Mathematica (Wolfram).

-For scoring invasion of the TFP1-iKD control and silenced parasites in HFF cells, samples were automatically scanned at a magnification of × 20 under an OlympusScan^R automated inverted microscope (3 wells per condition, 16 fields of acquisition per well). Image processing with the Cell^R software successively included signal-to-noise ratio optimization to allow cell nuclei segmentation, channel-associated image detection, and image subtraction (extracellular zoites subtracted from extra-plus intracellular tachyzoites), and intracellular tachyzoite segmentation using an edge detection algorithm as described in (Touquet et al., 2018).

### *T. gondii* tachyzoite surface flow analysis using microbead translocation assays

Two types of membrane flow assays were performed. The first provided a *posteriori* measure of membrane flow efficiency by quantifying the number of parasites for which microbeads were detected as clusters at their posterior pole after a 5 min time incubation with microbeads, and this under different adhesion and buffer conditions. The second provided information on the flow velocity profile by monitoring in real time the apicobasal displacement of microbeads attached to the parasite surface. In both cases, we used 0.2 μm diameter carboxylate-modified polystyrene fluorescent microspheres (FluoSpheres, far red/FR λ=660/680, 2% solid diluted in water, F8807, ThermoFisher Scientific) that were activated to promote covalent coupling with primary amines of proteins exposed on the parasite surface, thereby allowing bead attachment randomly on the parasite body. The activation was achieved using 50 μL of the initial microbead’s solution following the two-step 1-ethyl-3-[3-dimethylaminopropyl] carbodiimide (EDC)/sulfo NHS covalent coupling (Estapor carboxyl-modified dyed microspheres protocol, Merck Millipore) in MES buffer (30 min, room temperature) prior to extensive washing and recovery in 50 μL of the same buffer. The activated microbeads were sonicated (water bath Sonorex super RK31, Bandelin) for at least 10 min to break apart bead clusters immediately before the flow assays. Alternatively, they were coupled with 1 mg/mL of anti-SAG1 antibodies (NCL-Tg, Leica).

The freshly egressed or intracellular Δ*ku80* tachyzoites (WT, Δ*myoA,* or TFP1-iKD) were prepared as described above. For each condition tested (ICB *vs* MB, fibronectin *vs* PLL-PEG, addition of actin inhibitor or microneme secretagogue), a 100 μL of parasite suspension was incubated with 1 µL of activated microbeads for 3 min. The suspension was then deposited on either fibronectin or PLL-PEG coated coverslips inside a 4-well plate and gently settled (300g, 2 min) prior to be incubated at 37°C and 5% CO_2_ for 3 min. The microbead translocation assay was stopped by fixing the cells by addition of 3.2% PFA in the buffer. Because the PFA-fixed parasites were loosely sedimented on the non-adhesive PLL-PEG substrate and would be lost during the IFA procedure, we transferred them onto a PLL-coated coverslip and promoted cell attachment by centrifugation (300g, 3 min). For the assays requiring to inhibit actin dynamics, the tachyzoites were pre-incubated 5 min in MB supplemented with 1 μM cytochalasin D (C8273, Sigma) prior to addition of microbeads. To boost microneme exocytosis, the secretagogue zaprinast (Z0878, Sigma) was added (100 μM) together with the microbeads to the tachyzoite suspension. Membrane flow was also assessed in tachyzoites, which were left in suspension in MB in Eppendorf tubes under continuous mechanical rotation (15 rpm), followed by fixing on PLL-coated coverslips. All samples were immunostained to define the parasite contours as described in the “Lipid membrane staining and immunostaining” section. Labeled samples were rapidly imaged under confocal or widefield microscopes by collecting 0.3 μm z-image stacks from 20 to 30 fields of view in duplicates and for a minimum of 3 independent replicates. Parasites free of microbeads were excluded from the analysis while those associated with microbeads were enumerated as bearing beads on the body surface or specifically at the posterior pole. In the latter situation, microbeads were usually visualized as clusters which resulted from the apicobasal translocation of several microbeads.

For fast real time monitoring of microbead translocation on parasites settled either on fibronectin or PLL-PEG surfaces, 50 μL of Δ*ku80* WT tachyzoites suspension in ICB was incubated with 1 µL of activated carboxylated microbeads and deposited in the centre of a 35 mm diameter MatTeck chamber (Life Sciences). When cells have sedimented, the focus was adjusted and the ICB drop was carefully diluted in excess of MB (>2 mL). Videorecording was started as soon as the tachyzoites displayed conoid extrusion in response to the calcium containing MB as specified in the “dynamic and static microscopy” section (microbead/membrane flow fast monitoring details).

### Ultrastructure Expansion microscopy (U-ExM)

For U-ExM on micropatterned surfaces, 12 mm diameter glass coverslips were micropatterned with PLL-PEG as described above. The fibronectin conjugate displaying the Atto^488^ fluorophore and characterized by high quantum yield and photo-stability, was found to better resist the harsh denaturing conditions of the U-ExM protocol than the other conjugates. It allowed us to unambiguously delineate the micropatterns on expanded samples while their initial fixed dimension (2, 4, 6, 8 μm) provided direct control for the correct xy isotropic specimen enlargement. To produce the fibronectin-Atto^488^ conjugate, we incubated hydrophilic Atto^488^-NHS ester (41698, Sigma Aldrich) with 1 mg of fibronectin (1 mg/mL) at a 10:1 molar ratio in carbonate buffer (1 h, room temperature). The conjugate was purified using a Zeba Spin Desalting columns (898882, Thermo scientific) and filtered through a 0.2 μm syringe filter prior to be aliquoted and stored at 4°C. Micropatterned coverslips were incubated with 25 μg/mL of the fibronectin-Atto^488^ conjugate in carbonate buffer for 30 min at room temperature. The fibronectin-Atto^488^/PLL-PEG micropatterned coverslips were then equilibrated in MB, placed on a 4 well plate in 400 μL of MB to which 100 μL of the parasite suspension was added dropwise. The plate was centrifuged at 300g for 1 min to synchronize cell sedimentation and parasites were then left to glide at 37°C for 2 min. In the case of the micropattern samples, to better preserve the parasite shape, a pre-fixation was performed by adding 3.6% formaldehyde for 5 min (FA, Sigma Aldrich) and was followed by 1 min exposure to chilled methanol (−20°C) and an overnight incubation at 37 °C in 0.7% FA and 1% acrylamide (01697, Sigma Aldrich) mixture in PBS. In the case of MIC2 secretion assay, tachyzoites were collected and deposited in IBC on either fibronectin or PLL-PEG surfaces and then exposed to MB for 3 min prior to a pre-fixation (3.6% formaldehyde for 5 min) and fixation (0.7% FA and 1% acrylamide, 37°C, overnight) steps. We next followed the Gambarotto protocol (Gambarotto et al., 2019) for sample gelation (1 h, 37°C), gel denaturation (1 h, 95 °C) and gel expansion in water (3 times 30 min baths). After incubation in PBS to shrink the gel, several ∼4 mm diameter pieces were excised and processed for immunostaining (see next section). The labelled gels were re-expanded with three 20 min water baths and immobilized on a PLL-coated glass coverslip in the microscope chamber to avoid gel drift during image acquisition. The 0.2 μm z-image stacks were captured with the Live SR super resolution module controlled by the 3D Live SR software (Gataca Systems). For the U-ExM tachyzoite samples on micropatterns, image stacks were resliced with Fiji prior to 3D reconstruction with 3D viewer plugin (Schmid et al., 2010) from which movies were further retrieved. For the MIC2 stained U-ExM-expanded tachyzoite samples, a region of interest was defined as the last fifth part of the tachyzoite body, measured from the long axis of the cell on the image that represents the maximal intensity projection from the image stack. Pixel intensity values of the region of interest were retrieved using Fiji.

### Lipid membrane staining and immunostaining of fixed samples

-Lipid membrane staining: live tachyzoites prepared as mentioned above were incubated with the lipid dye PKH26 to label the plasma membrane using the Cell Linker kit (041M0852, Sigma Aldrich) in 0.5 mL final volume following the manufacturer protocol. Labeled parasites were resuspended in MB and captured by live confocal microscopy using the λ=561 nm laser and the x60 objective (20 fields of view per assay, 2 independent assays). Image stacks were processed to obtain an intensity maximal projection image from which the length of the parasites along the major axis was measured using the Fiji software.
-Immunostaining of fixed tachyzoite samples for quantitative bead flow analysis, TF1 cell line characterization: PFA fixed samples on coverslips were successively incubated in 50 mM NH_4_Cl in PBS (10 min) to quench the free aldehyde groups and in BSA (2% w/v, Sigma)-containing PBS to prevent non-specific binding of the subsequently used antibodies. For all the membrane flow analysis, the parasite contours were detected by immunostaining of the P30 surface-exposed protein. To control the efficiency of TFP1 silencing in the TFP1-iKD cell line, vehicle (ethanol) and ATc-treated free parasites were first permeabilized using Triton X-100 (0.1% v/v, Sigma) for 5 min prior to the blocking and staining steps using antibodies against the glideosome-associated protein GAP45 and anti Ty tag antibodies in successive steps. In addition, tachyzoite invasiveness on HFF was scored by labeling the P30 protein (targeting only extracellular parasites), followed by TX100 permeabilization and GAP45 staining (targeting both extra and intracellular parasites). Hoechst 33242 was added to label all nuclei (parasites and host cells). After washing in PBS and water, coverslips were mounted in Mowiol medium (475904, Merck Millipore).
-Immunostaining of the U-ExM processed samples: gel pieces were incubated with primary antibodies overnight at 4°C in 2% BSA in PBS prior to be washed in PBS-Tween 0.1 % and incubated with the secondary antibodies in 2% BSA in PBS for 3 h at 37 °C followed by PBS-Tween 0.1 % washes.
-For all staining procedures, first and secondary antibodies were used as described in the Supplementary Table 1. Images were collected and processed as described in each specific section.

### Functionalization of coverslips with PLL-PEG-b/SAv:b-X model coatings

18 mm glass coverslips were cleaned in 0.5 M HCl (overnight, 65°C) and extensively washed in water and ethanol prior to be used. The coverslips were treated with UV-ozone for 20 min and air gun-dried. The coverslips were first exposed to 500 μL of 0.1 mg/mL PLL-PEG-b in HEPES-NaCl buffer for 1 h, followed by rinsing with HEPES-NaCl buffer 5 times. Next, they were exposed to the streptavidin:biotinylated ligand (SAv:b-X) complexes (b-PEG derivatives, b-HS, b-HSA) in HEPES-NaCl buffer. The SAv:b-X mixtures were prepared in HEPES-NaCl buffer. SAv concentration was fixed to 10 µg/mL. The concentration of b-X ligands varied according to the desired SAv:b-X stoichiometry (1:0, 1:1, 1:2 or 1:3). The mixture was incubated for 15 min and then placed on the top of the PLL-PEG-b-functionalized glass coverslips. After 1h incubation, the layers were rinsed with HEPES-NaCl buffer 5 times and with MB, in which they were kept until motility studies.

### Quartz crystal microbalance with dissipation monitoring (QCM-D) measurements and viscoelastic modeling

QCM-D measurements were performed using a Q-Sense E4 system equipped with four Q-Sense flow modules (Biolin Scientific). The measurements were done in a flow mode (flow rate = 20-50 μL/min) at *T* = 24 °C. Silica-coated quartz sensors (Biolin Scientific) were used as substrates. The sensors were cleaned using UV-ozone for 20 min, followed by rinsing with water and ethanol and air gun-dried before installing them into the QCM-D modules. The normalized frequency (Δ*f =* Δ*f*_j_/j) and dissipation (Δ*D*) shifts were recorded for six overtones (*j* = 3, 5, 7, 9, 11, and 13), in addition to the fundamental resonance frequency (4.95 MHz), using the QTools software. Changes in Δ*f* and Δ*D* for j = 7 are presented, while other overtones showing similar trends.

For PLL-PEG, after recording a baseline in HEPES-NaCl buffer, 0.1 mg/mL PLL-PEG was injected in HEPES-NaCl buffer and rinsed using the same buffer. For PLL, after recording baseline in water, 0.1% w/v PLL was injected in water and then rinsed with water. For fibronectin, after recording a baseline in PBS, 50 µg/ml fibronectin was injected in PBS and rinsed with the same buffer. For the tunable surfaces based on SAv chemistry, after recording a baseline in HEPES-NaCl buffer, 0.1 mg/mL PLL-PEG-b was injected in HEPES-NaCl buffer and rinsed in the same buffer. After that, SAv:b-X mixtures were prepared at desired stoichiometry (at SAv concentration fixed to 10 µg/mL), incubated 15 min and injected until signal stabilization, followed by rinsing with HEPES-NaCl buffer. The thickness (*d*) and viscoelastic properties of the adsorbed layers were determined by fitting the QCM-D data to a continuum viscoelastic model (Johannsmann, 1999) implemented in the QTM software (D. Johannsmann, Technical University of Clausthal, Clausthal-Zellerfeld, Germany). The fitting procedure was described in detail previously (Eisele et al., 2012). Viscoelastic properties were parameterized in terms of the shear storage modulus *G*’(*f*) and the shear loss modulus *G*’’(*f*). The frequency dependencies of the storage and loss moduli were assumed to follow power laws within the measured range of 15 to 65 MHz, with exponents α’ and α’’, such that *G*(*f*) = *G*_0_ (*f*/*f*_0_)^α^, with *f*_0_ set to 15 MHz. The data were fitted using a small load approximation (SLA),-at a fundamental frequency of 4.95 MHz. The bulk properties of the water or buffer solution were fixed at density ρ=1.0 g/cm^3^ and viscosity η=0.89 mPa. The density of the hydrated film was assumed to be 1.0 g/cm^3^.

### Statistical analysis

All the motility and membrane flow-related data were statistically analyzed using GraphPad Prism 8.0 software for Windows (La Jolla, CA, USA). Plots were made using the same software, and data are presented as mean ± standard deviation if not indicated otherwise, and exported to CorelDraw Graphics Suite 2017 (Alludo) for visualization purposes. Statistical tests and exact p-values are provided in the corresponding captions, and a comprehensive list is shown in the Supplementary Table 2.

