## Extended Figures and Supplemental Tables for "Uncovering the simple adhesive strategy of the Toxoplasma parasite for high-speed motility"

Extended Figure 1 - Vigetti et al.

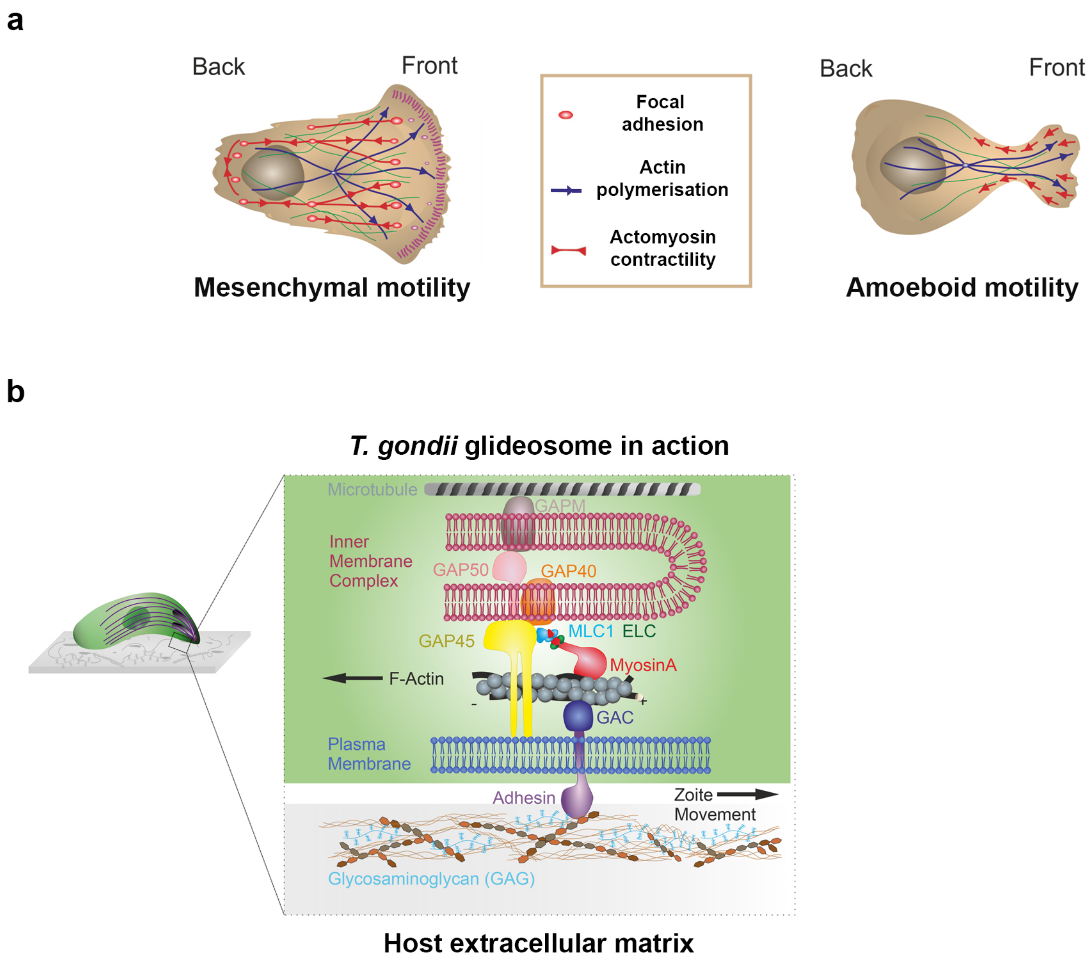


**Schematics of crawling motile strategies used by eukaryotic cells and *T. gondii* molecular motile machinery. (a)** Schematic of non-swimming eukaryotic cells undergoing distinct motility modes, grouped as crawling (mesenchymal) or bleb-like (amoeboid) motility. Mesenchymal motility relies on the formation of a flat lamellipodium at the cell front, driven by actin filament polymerization (blue line) and coupled to the substrate through the assembly of dynamic focal adhesions (red circles), which, once mature, support force transmission and traction while actomyosin contractility occurs at the cell rear. Instead, amoeboid motility proceeds without mature focal adhesions, hence without traction, and relies on the contractility of the actomyosin cortex coupled to the plasma membrane to promote bleb-like deformations and forward displacement. **(b)** Schematic of the T. gondii glideosome machinery components, centered on the sub-membranous myosin A motor and regulatory units distributed between the parasite plasma membrane and the beneath inner membrane complex (IMC) through the contribution of glideosome associated proteins (GAPs). Actin filaments are nucleated apically and pulled backward by the myosin motors in a fixed orientation, thereby proposed to generate the mechanical force powering forward motion.

Extended Figure 2 - Vigetti et al.

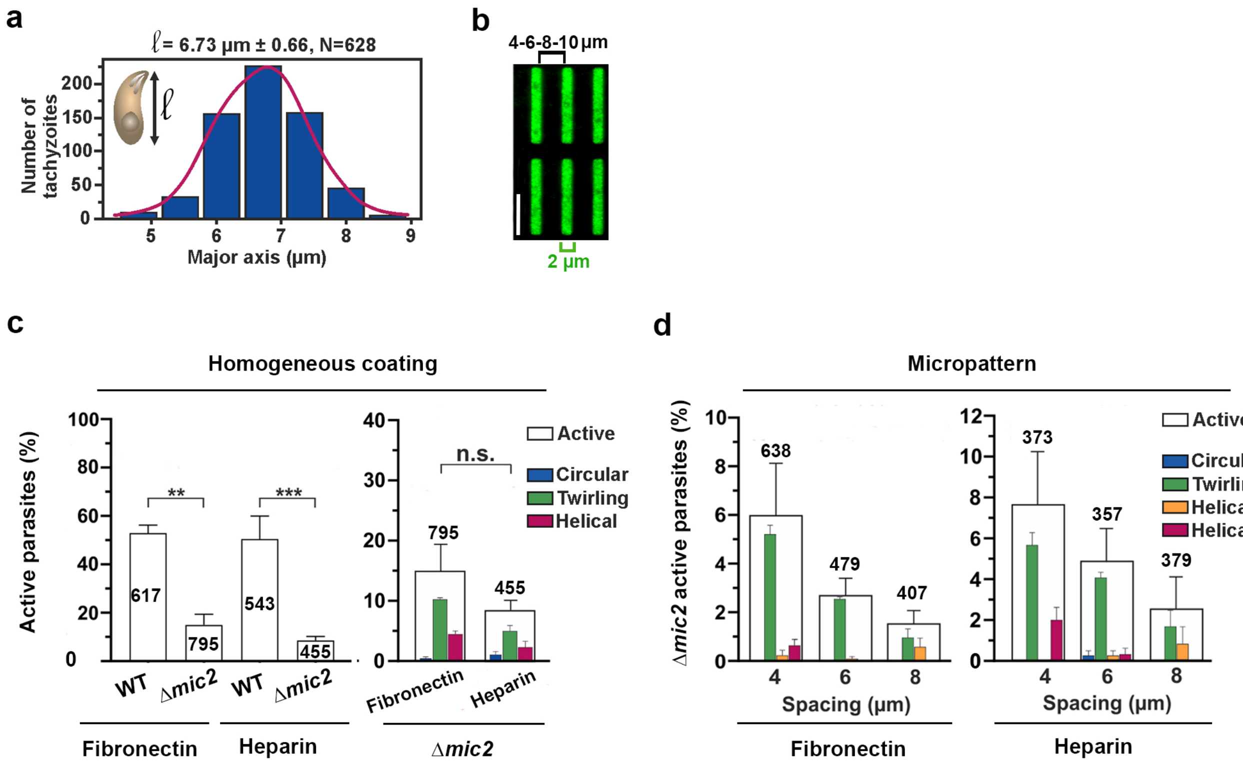


**Micropattern design and Δmic2 tachyzoite motile behavior analysis on homogeneous and micropatterned surfaces. (a)** Histogram for the major axis length of a tachyzoite population (628 cells) measured from 2 independent videorecording sessions after labeling the parasite surface with the fluorescent PHK26 lipophilic dye (λex 551 nm; λem 567 nm). **(b)** Confocal image of the micropatterns coated with a mixture of fibronectin and fluorescent fibronectin-HiLyte488 which have been designed as pro-adhesive rectangular micropatterns (2x20 µm) spaced by anti-adhesive PLL-PEG lines of increasing width as indicated (4 µm shown), based on the parasite length distribution shown in **(a)**. Scale bar: 10 µm. **(c)** Percentages of active WT and Δmic2 tachyzoites on fibronectin and heparin homogeneous coatings. On the right graph, the active Δmic2 tachyzoites are subcategorized according to the motile behaviors with color codes (circular, twirling and helical) on fibronectin and heparin substrates (n = 455 to 795 parasites from at least 3 experiments). **(d)** Percentages of active Δmic2 tachyzoites on fibronectin and heparin micropatterns spaced by PLL-PEG lines of either 4, 6 or 8 µm width. The active Δmic2 tachyzoites are subcategorized according to the three motile behaviors (n = 357 to 638 parasites from at least 3 experiments). Statistical analyses are detailed in Materials and Methods section.

Extended Figure 3 - Vigetti et al.

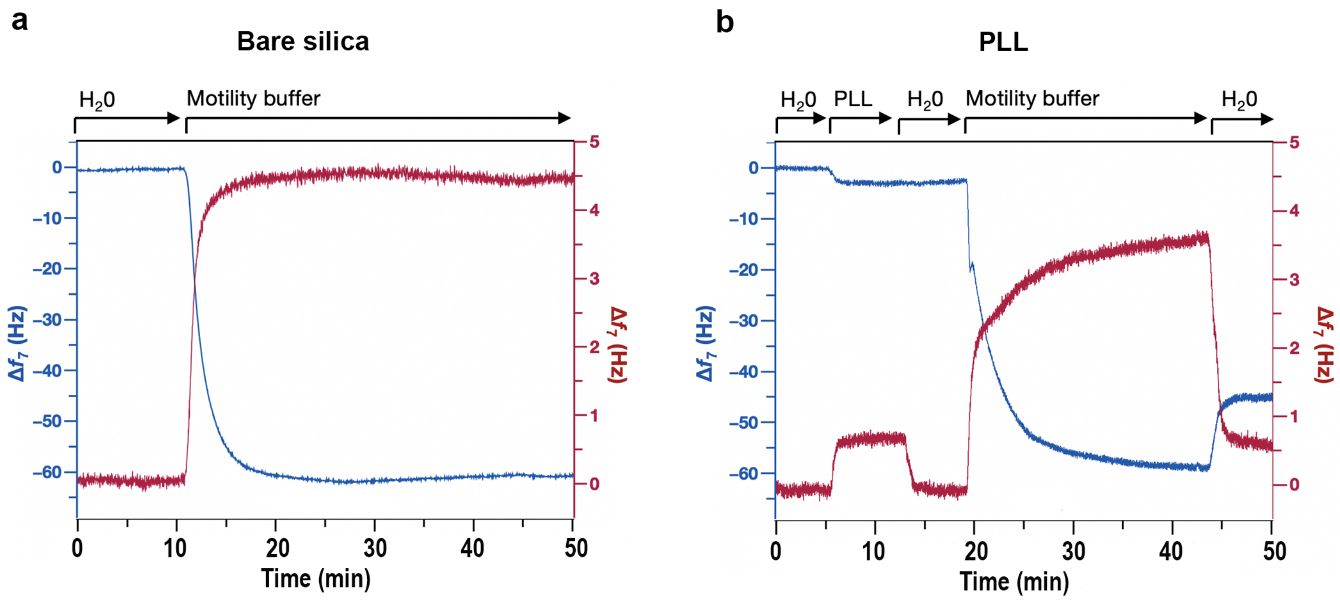


**QCM-D characterization of bare and PLL-coated silica surfaces exposed to the tachyzoite motility buffer. (a)** QCM-D shows the presence of non-specific adsorption of the components of the tachyzoite motility buffer (HBSS buffer with fetal calf serum and CaCl2) to the bare silica surface, evidenced by the strong frequency and dissipation shift upon their injection. **(b)** QCM-D shows fast binding of PLL to the silica surface, resulting in a rigid ($\Delta D \approx 0$) coating, stable in water. The components of the tachyzoite motility buffer show strong attachment to the PLL layer, evidenced by the strong $\Delta f$ and $\Delta D$ shifts upon their injection. The binding of the motility buffer components is only partially reversible as evidenced by the residual QCM-D shifts upon water rinsing.

Extended Figure 4, Vigetti et al.

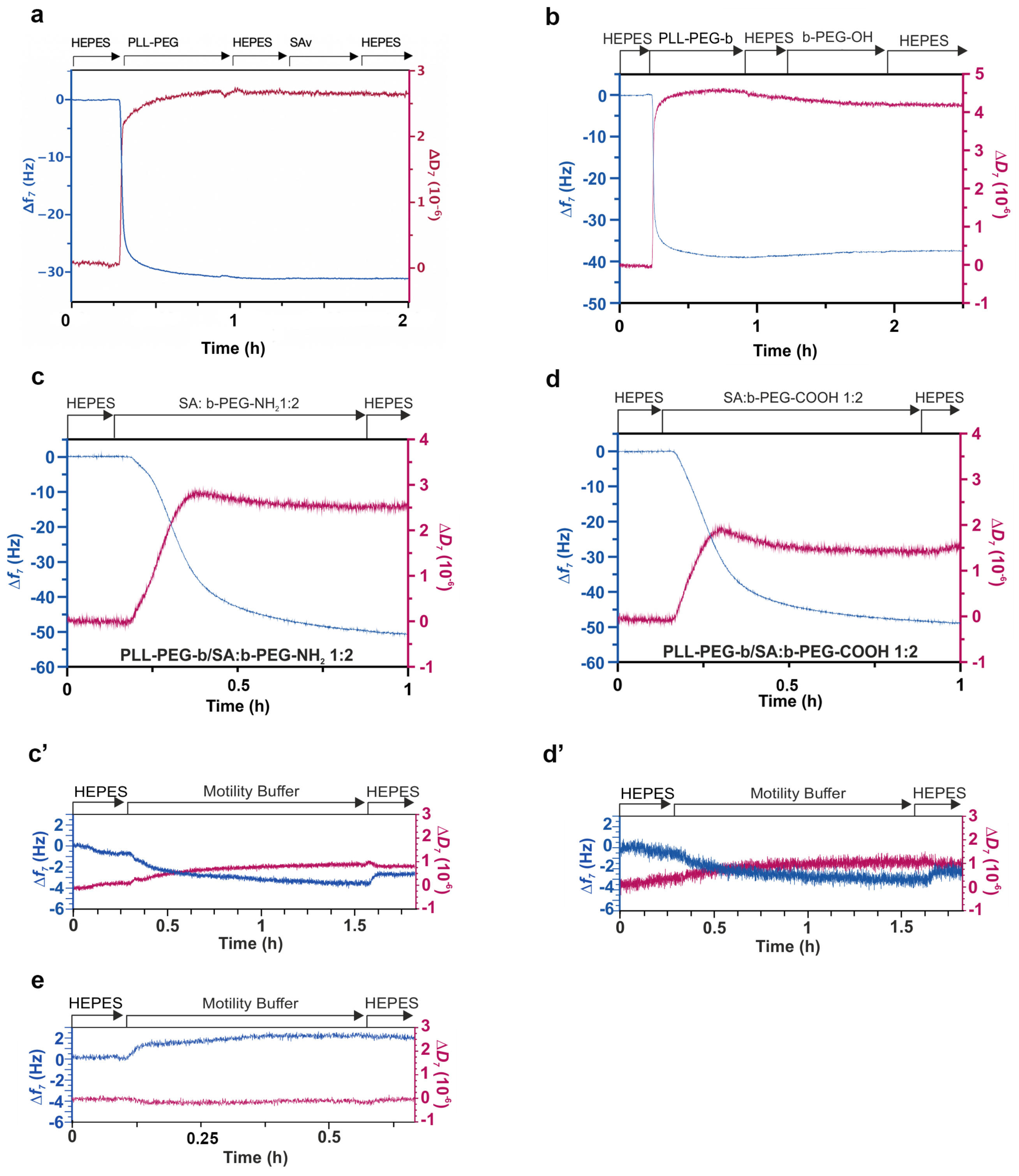

**QCM-D confirms the absence of non-specific interactions and specific binding to model surfaces based on PLL-PEG and SAv chemistry.** **(a)** QCM-D shows the absence of SAv (c = 10 µg/mL) binding to the PLL-PEG layer lacking biotins. **(b)** QCM-D shows no binding of b-PEG-OH (c = 100 µg/mL) to PLL-PEG-b in absence of SAv interlayer. **(c-d)** QCM-D monitoring of binding of SAv:b-PEG-NH2 (1:2) **(c)** and SAv:b-PEG-COOH (1:2) **(d)** complexes to the PLL-PEG-b layer resulting in a stable and soft ($\Delta D > 0$) coatings. **(c', d', e)** The components of the tachyzoite motility buffer (i.e., glucose, fetal calf serum) do not show attachment to the formed PLL-PEG-b/SAv:b-PEG-NH2 (1:2) **(c')**, PLL-PEG-b/SAv:b-PEG-COOH (1:2) **(d')** and PLL-PEG-b/SAv:b-HSA (1:1) **(e)** coatings, evidenced by the absence of strong $\Delta f$ and $\Delta D$ shifts upon their injection and rinsing; for QCM-D monitoring of PLL-PEG-b/SAv:b-HSA formation, see Fig. 5c in the manuscript (circles).

Extended Figure 5 - Vigetti et al.

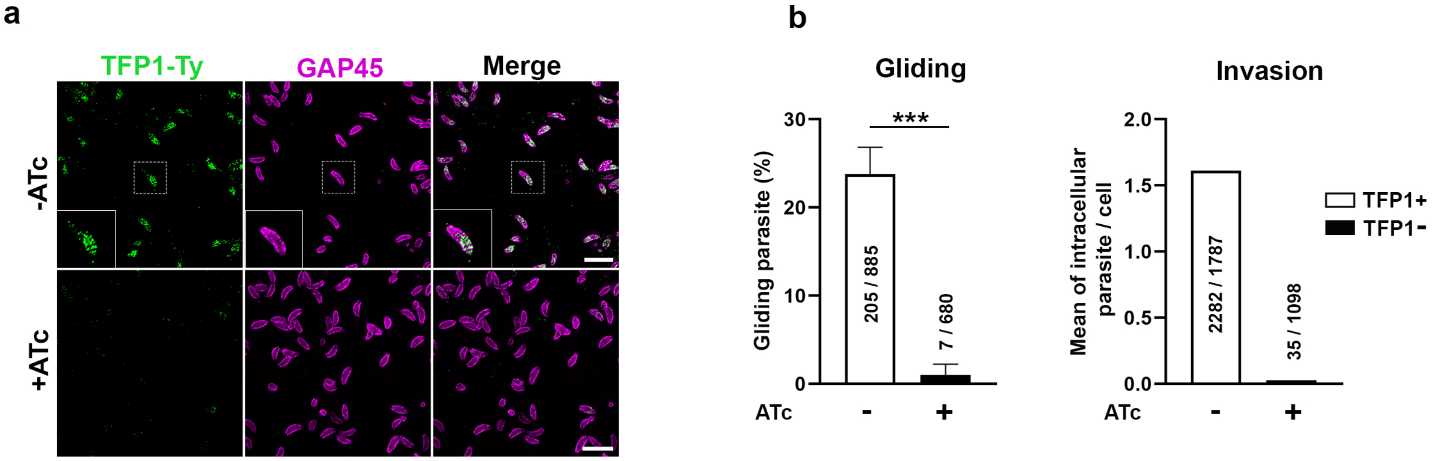


**Characterization of T. gondii tachyzoite conditionally silenced for TF1 expression. (a)** Confocal images of the Tati-TFP1-3xTy tachyzoites freshly egressed from HFF cells exposed ~ 48 h to anhydrotetracycline (1 µg/mL) (+ ATc, bottom row) or to the 100% ethanol vehicle (- ATc, top row). Parasite contours are delineated with GAP45 immunostaining (magenta) and TFP1 is immunodetected with anti-Ty tag antibodies (green). Scale bar: 5 µm. **(b)** Graph showing (left) the percentage of gliding cells for a 5 min video-sequence, and (right) the average number of intracellular tachyzoites per HFF cell after 15 min of invasion for the Tati-TFP1-3xTy tachyzoites expressing (ATC -) or depleted in TFP1 (ATc +). Both assays represent triplicates from one representative experiment.

### Supplemental Table 1

| Antibody | Application | Handling | Source |
|---|---|---|---|
| Mouse α-TgP30<br>Clone TP3 | IF | 1:600, 20 min, 23°C | Novocastra (Leica)<br>Ref: NCL-TG; batch: 564038 |
| Mouse α-Ty | IF | 1:1000, 1h, 23°C | D. Soldati-Favre<br>(University of Geneva) |
| Rabbit anti-TgGAP45 | IF<br>U-ExM | 1:2000, 1h, 23°C<br>1:500, ON, 4°C | D. Soldati-Favre<br>(University of Geneva) |
| Rat anti-TgMIC2<br>Clone 11 | U-ExM | 1:100, ON, 4°C | D. Sibley |
| Mouse anti-α-tubulin<br>mAb clone DM1A | U-ExM | 1:400, ON, 4°C | Millipore<br>Ref: 05-829; batch:3925603 |
| Rat anti-HA (IgG1)<br>mAb clone 3F10 | U-ExM | 1:200, ON, 4°C | Roche<br>Ref: 12158167001; batch: |
| Goat anti-mouse (IgG H+L)-<br>Alexa Fluor™ 488 (HCA) | IF | 1:600, 30 min to 1h, 23°C | Invitrogen<br>Ref: A11029; batch: 1073083 |
| Goat anti-mouse (IgG H+L)-<br>Alexa Fluor™ 594 (HCA) | IF | 1:600, 1h, 23°C | Invitrogen<br>Ref: A11032; batch: 2301112 |
| Goat anti-rabbit (IgG H+L)-<br>Alexa Fluor™ 488 (HCA) | IF<br>U-ExM | 1:600, 1h, 23°C<br>1:400, 3 h, 37°C | Invitrogen<br>Ref: A11034, batch : 1073084 |
| Goat anti-rat (IgG H+L)-<br>Alexa Fluor™ 488 (HCA) | U-ExM | 1: 400, 3h, 37°C | Invitrogen<br>Ref : A21208, batch : 20922 |
| Goat α-mouse IgG STAR RED | U-ExM | 1:200, 3h, 37°C | Abberior<br>Ref: STRED-1001,<br>Batch: 10601PK-4 |

| | | | |
|---|---|---|---|
| Goat anti-rat (IgG H+L)-Alexa Fluor$^{TM}$ 568 (HCA) | U-ExM | 1: 400, 3h, 37°C | Invitrogen<br><br>Ref : A11036, |

**Supplementary Table 2 : Statistical tests and exact p-values for all motility and membrane flow assays** (NS: non-significant, p-value >0.05; *: p-value <0.05; **: p-value <0.01; ***: p-value <0.001; ****: p-value <0.0001.)

| **Figure** | **Comparison** | **Test** | **p-value** |
|---|---|---|---|
| 1b | Adhesion on Heparin *vs* fibronectin | Unpaired t test | NS, p=0.5647 |
| 1b | Adhesion on PLL-PEG *vs* fibronectin | Unpaired t test | **, p=0.0024 |
| 1b | Adhesion on PLL-PEG *vs* heparin | Unpaired t test | **, p=0.0015 |
| 2c | Activity on fibronectin full coat *vs* fibronectin micropatterned surfaces | One-way ANOVA followed by Dunnet's multiple comparisons against fibronectin full coat (control) | NS, p=0.2015<br>Dunnet's comparisons:<br>MP 4µm: p=0.6815<br>MP 6 µm: p= 0.9983<br>MP 8µm: p=0.1261<br>MP 10µm: p=0.2930 |
| 2e | Activity on Fibronectin full coat *vs* PLL-PEG | Unpaired t test | ****, p<0.0001 |
| 2d | Activity on Heparin full coat *vs* heparin micropatterned surfaces | ANOVA followed by Dunnet's multiple comparisons against heparin full coat (control) | NS, p=0.7767<br>Dunnet's comparisons:<br>MP 4µm: p=0.9975<br>MP 6 µm: p= 0.7484<br>MP 8µm: p=0.9773 |
| 2f | Activity on Heparin full coat *vs* PLL-PEG | Unpaired t test | ***, p=0.0005 |
| 2f | Activity on Heparin full coat *vs* Heparin-FITC full coat | Unpaired t test | NS, p=0.4951 |

| | | | |
|---|---|---|---|
| 2g | Total trajectory of helical gliding across micropatterns on fibronectin full coat *vs* fibronectin micropattern | One-way ANOVA followed by Dunnet's multiple comparisons against fibronectin full coat | NS, p=0.1169<br>Dunnet's comparisons:<br>MP 4µm: p=0.2560<br>MP 6 µm: p=0.5763<br>MP 8µm: p=0.9877 |
| 2g | Mean velocity of helical gliding across micropatterns on fibronectin full coat *vs* fibronectin micropattern | One-way ANOVA followed by Dunnet's multiple comparisons against fibronectin full coat | ****, p<0.0001<br>Dunnet's comparisons:<br>MP 4µm: **, p=0.0018<br>MP 6 µm: ns, p=0.0847<br>MP 8µm: ****, p<0.0001 |
| 2g | Total trajectory of helical gliding across micropatterns on heparin full coat *vs* heparin micropattern | One-way ANOVA followed by Dunnet's multiple comparisons against heparin full coat | NS, p=0.8114<br>Dunnet's comparisons:<br>MP 4µm: ns, p=0.9953<br>MP 6 µm: ns, p=0.9231<br>MP 8µm: ns, p=0.8361 |
| 2g | Mean velocity of helical gliding across micropatterns on heparin full coat *vs* heparin micropattern | One-way ANOVA followed by Dunnet's multiple comparisons against heparin full coat | NS, p=0.4282<br>Dunnet's comparisons:<br>MP 4µm: ns, p=0.9577<br>MP 6 µm: ns, p=0.9844<br>MP 8µm: ns, p=0.2569 |

| | | | |
|---|---|---|---|
| 6a | Activity on PLL-PEG *vs* PLL-PEG-b | Mann Whitney test | NS, p=0.1333 |
| 6a | Activity on PLL-PEG *vs* PLL-PEG-b/SA | Mann Whitney test | NS, p=0.1333 |
| 6c | Activity on PLL-PEG *vs* PLL-PEG-b/SA-PEG-OH 1:1 | Mann Whitney test | NS, p=0.6571 |
| 6c | Activity on PLL-PEG *vs* PLL-PEG-b/SA-PEG-NH2 1:2 | Fisher's exact test | ****, p<0.0001 |
| 6c | Activity on PLL-PEG-b/SA-PEG-NH2 1:2 *vs* PLL-PEG-b/SA-PEG-NH2 1:3 | Fisher's exact test | NS, p=0.4687 |
| 6c | Activity on PLL-PEG *vs* PLL-PEG-b/SA-PEG-COOH 1:2 | Fisher's exact test | ****, p<0.0001 |
| 6c | Activity on PLL-PEG-b/SA-PEG-COOH 1:2 *vs* PLL-PEG-b/SA-PEG-COOH 1:3 | Fisher's exact test | **, p=0.0091 |
| 6b | Activity on PLL-PEG *vs* PLL-PEG-b/SA-PEG-HSA 1:1 | Fisher's exact test | ****, p<0.0001 |
| 6b | Activity on PLL-PEG-b/SA-PEG-HSA 1:1 *vs* PLL-PEG-b/SA-PEG-HSA 1:3 | Fisher's exact test | *, p=0.0306 |
| 6d | Activity on PLL-PEG *vs* PLL-PEG-b/SA-PEG-HS 1:1 | Fisher's exact test | ****, p<0.0001 |
| 6d | Activity on PLL-PEG-b/SA-PEG-HS 1:1 *vs* PLL-PEG-b/SA-PEG-HS 1:2 | Fisher's exact test | **, p=0.0055 |
| 6e | Activity on Heparan sulfate treated with Surfen 0 µM *vs* 4 µM | Unpaired t test | **, p=0.0030 |
| 6e | Activity on Heparan sulfate treated with Surfen 0 µM *vs* 20 µM | Unpaired t test | *, p=0.0176 |
| 6e | Activity on Heparan sulfate treated with Surfen 4 µM *vs* 20 µM | Unpaired t test | NS, p=0.1804 |
| 7d | Tachyzoites with posterior beads, Motility Buffer control *vs* Cyto D+, Fibronectin surface | Unpaired t test | ***, p=0.0005 |
| 7d | Tachyzoites with posterior beads, Motility Buffer control *vs* ICB, Fibronectin surface | Unpaired t test | ***, p=0.0004 |

| | | | |
|---|---|---|---|
| 7d | Tachyzoites with posterior beads, Motility Buffer control *vs* Zaprinast +, Fibronectin surface | Unpaired t test | NS, p=0.2139 |
| 7d | Tachyzoites with posterior beads, Motility Buffer control *vs* Cyto D +, PLL-PEG surface | Unpaired t test | ****, p<0.0001 |
| 7d | Tachyzoites with posterior beads, Motility Buffer control *vs* ICB, PLL-PEG surface | Unpaired t test | ****, p<0.0001 |
| 7d | Tachyzoites with posterior beads, Motility Buffer control *vs* Zaprinast +, PLL-PEG surface | Unpaired t test | *, p=0.0406 |
| 7e | Tachyzoites with posterior beads, *Δmyoa*, fibronectin vs PLL-PEG surfaces | Unpaired t test | NS, p=0.4628 |
| 7g | Tachyzoites with posterior beads, Motility Buffer TFP1+ *vs* TFP1- | Unpaired t test | NS, p=0.7000 |
| 7g | Tachyzoites with posterior beads, Motility Buffer+Zaprinast on TFP1+ *vs* Motility Buffer+Zaprinast TFP1- | Unpaired t test | NS, p=0.8000 |
| 7h | MIC2 basal restricted fluorescence, fibronectin surface, ICB *vs* MB, | Unpaired t test | p=0.0029 |
| 7h | MIC2 basal restricted fluorescence, ICB on fibronectin surface *vs* MB on PLL-PEG surface | Unpaired t test | p<0.0001 |
| 7h | MIC2 basal restricted fluorescence, MB, PLL-PEG *vs* Fibronectin surfaces | Unpaired t test | p=0.0081 |
| Ext2c | Active parasites, fibronectin, WT *vs* *Δmic2* | Unpaired t test | **, p=0.0016 |
| Ext2c | Active parasites, heparin, WT *vs* *Δmic2* | Unpaired t test | ***, p=0.001 |
| Ext2c | Active parasites, *Δmic2,* fibronectin *vs* heparin | Unpaired t test | ns, p=0.1256 |
| Ext5b | Gliding parasite | Fisher’s exact test | ***, p=0.0004 |
